# ENHANCED EFFICIENCY OF RNA-GUIDED CAS12a VERSUS CAS9 TRANSGENE KNOCK-IN AND ACTIVITY AT A *SCHISTOSOMA MANSONI* GENOME SAFE HARBOR

**DOI:** 10.1101/2023.09.12.557428

**Authors:** Max F. Moescheid, Prapakorn Wisitphongpun, Victoria H. Mann, Thomas Quack, Christoph Grunau, Christoph G. Grevelding, Wannaporn Ittiprasert, Paul J. Brindley

## Abstract

Recently, we reported programmed Cas9 mediated insertion of a reporter gene into a gene safe harbor site, GSH1, of *Schistosoma mansoni* via homology-directed repair (HDR) using overlapping guide RNAs. Here, we report efficient and precise CRISPR/Cas12a-mediated homology directed insertion (knockin, KI) of a 5’ C6-PEG10-modified double-stranded transgene bearing microhomology arms, 50 nt in length, at GSH1. At the outset, we undertook bioinformatic and computational analysis following by experimental verification of the regulatory activity of endogenous schistosome ubiquitin (SmUbi) promoter and terminator, to drive strong reporter gene expression. Green fluorescent protein activity driven by SmUbi followed electroporation-mediated transfection of schistosome eggs. HDR induced by RNA-guided CRISPR/Cas12a, which releases overhanging DNA strands of 18-24, delivered more efficient KI than CRISPR/Cas9. In this non-model pathogen, programmed KI facilitated precise chromosomal integration of the reporter-gene with at GSH1. The approach advances schistosome transgenesis field and may also advance functional genomics and transfection methods in related parasitic and non-parasitic helminths, which hitherto lack these tools.

**Author summary:** Genome editing (CRISPR) technology is revolutionizing advances in biology, medicine, and agriculture. Transgenesis approaches are integral in diverse applications including gene therapy, biotherapeutics, deciphering host-pathogen interactions, and enhancements in agricultural production. Parasitic worms that are responsible for infectious diseases including neglected tropical diseases (NTDs), which cause substantial morbidity and mortality. NTDs mainly occur in the Global South, and they are responsible for a disease burden that exceeds that caused by malaria and tuberculosis. Infections with parasitic helminths also are responsible for immense economic burden in the agriculture. Tools for functional genomics in parasitic helminths are limited. Access to CRISPR-based approaches can be expected to hasten development of drug and/or vaccine targets for these diseases. Here, we focused on the helminth *Schistosoma mansoni*, a water borne parasite of humans, and which is endemic in Africa, and northeastern South America. To advance the state of the art in laboratory techniques currently used to study the biology and pathogenesis of this and related pathogens, we evaluated a spectrum of technological approaches aimed at improved current lab practice in this field. The findings demonstrated that specific technical and chemical modifications, including deploying a DNA cutting enzyme termed Cas12a along with a transgene with chemically modified short flanking sequences (homology arms) provided improved gene editing efficiency for this schistosome.

## INTRODUCTION

*Schistosoma mansoni*, a blood-dwelling parasite of humans and animals, is the cause of the infectious disease schistosomiasis. As with all trematodes, *S. mansoni* and related species exhibit complex life cycles including an invertebrate snail host and a vertebrate final host. Schistosomiasis treatment relies on a single drug, praziquantel (PZQ) (1). Due to the fear of upcoming PZQ resistance (2), there is the urgent need for the development of alternative treatment strategies. For this, and to understand parasite biology as such, it is inevitable to characterize the function of target genes and their potential role(s) in, for example, parasite development and homeostasis.

Substantial effort has been exerted to decipher the *S. mansoni* genome (3–5). In addition, ongoing efforts have aimed at deciphering the *S. mansoni* transcriptome of the adult as well further developmental stages (6–16). Now, in the rising post-genomic/transcriptomic era of *S. mansoni* research, new tools are needed for the functional characterization of genes of interest (GOI). Until now, RNA interference (RNAi) has been proven as the most suitable method for functional gene characterization. However, RNAi efficiency varies, and it can lead to ectopic effects (17–20). To study GOI, knock-out (KO) models are common for various model organisms, but not yet established for trematodes or other helminths. Towards this end, a recent breakthrough achieved CRISPR/Cas9 transgene KI into a schistosome genome safe harbor site (21).

CRISPR/Cas-mediated genome editing has revolutionized, or soon will, research in biomedicine and the life sciences generally (22, 23). An increasing repertoire of Cas homologues, in addition to Cas9, possessing divergent programmed nuclease activity including alternative PAMs, selectivity for diverse nucleic acid substrates, and sensitivity to metal ions, to temperature, and other characteristics have been reported (24, 25). Cas12a is enzymatically active in eggs of *S. mansoni* where a Cas12a orthologue from *Acidaminococcus* (AsCas12a) more efficiently edited the *omega-1* locus than Cas9 (26). CRISPR/Cas9 RNPs are assembled with two components – guide RNA (gRNA) and Cas9 nuclease (27–29). This nuclease induces blunt end-DSBs (double stranded breaks) within a programmed target, which can be repaired by genome maintenance mechanisms including non-homologous end joining (NHEJ) and homology-directed repair (HDR). CRISPR/Cas12a components consist of a crRNA and a Cas12a nuclease that releases overhanging strands at the programmed site (30). CRISPR/Cas9 recognizes the NGG protospacer adjacent motif (PAM) to catalyze the DSB upstream of PAM whereas CRISPR/Cas12a recognizes a T-rich PAM, TTTV, with the DSB downstream of the PAM (31). Both nucleases have utility for gene editing in discrete genomic contexts; Cas9 may have advantages for editing GC-rich regions, Cas12a may be better suited for editing AT-rich regions (32–34).

Transgene insertion in the genome can negatively impact the host cell function or the cell may silence the transgene due to chromatin structure. To surmount these potential problems, the gene therapy field developed the concept of the genome safe harbor (GSH) (35). An ideal GSH has been defined as a region that does not overlap functional genomic elements and that lacks heterochromatic marks that could impede transcription. We have predicted several GSHs in *S. mansoni* using criteria that included a location in euchromatin to avoid transgene silencing, a unique genome-target sequence to minimize off-target events, avoidance of long noncoding RNA-encoding genes, and the presence of histone marks for open chromatin structure and, conversely, the absence of epigenetic marks indicating heterochromatin (21). We reported the Cas9-mediated integration of an exogenous DNA sequence into GSH1 (genome safe harbor number 1) of *S. mansoni*. Programmed cleavage improved when three overlapping guide RNAs complexed with the Cas9 nuclease from *Streptococcus pyogenes* (21) were used in unison. As biological targets, we used liver-stage *S. mansoni* eggs (LE), which represent a heterogenous population at different stages of development. LE comprise newly laid eggs, intermediate, and mature eggs (from stages 1 through 8 of the staging system of Jurberg and coworkers (36)), and they were transfected with Cas9 or Cas12 ribonuclear protein complexes (RNPs). Gene knock-in (KI) was achieved in the presence of a donor repair template encoding enhanced green fluorescent protein (EGFP) bearing 5’-phosphorothioate-modified terminus HA of 300-600 bp in length. The transgene was 4,551 bp in length. The use of triple, overlapping gRNAs induced higher levels of precise transgene knock-in (KI) in comparison to single and double gRNAs. Nonetheless, further optimization aiming for simpler, less expensive, and enhanced levels of efficacy likely can be achieved.

To this end, here we evaluated the editing efficiencies of Cas9 versus Cas12a. In addition, we deployed a single gRNA in tandem with short, 5’-modified homology arms of the repair template, and finally investigated whether also newly laid eggs (NLE)| were amenable to transfection. This latter manipulation could ensure biologically standardized editing approaches compared to LE. We observed that both nucleases, Cas9 and Cas12a, induced on-target insertions and deletions (InDels). Notably, the editing efficiency of Cas12a significantly exceeded that of Cas9. Furthermore, with 5’-C6 amine-PEG10-modified donor template and minimal length HA of 50 bp flanking the donor transgene, we observed transgene KI at GSH1 as well as by the expression of the transgene EGFP. This hitherto unprecedented efficiency of transgene KI in schistosomes advance genome-editing approaches for *Schistosoma mansoni*.

## RESULTS

### Promoter elements of the schistosome ubiquitin gene

Access to a selectable marker gene under the control of regulatory elements, which are active in the desired way, would likely hasten the establishment of chromosomally modified schistosomes. Also desirable in some contexts is stage-independent reporter-gene activity. With these longer-term goals in mind, here we sought appropriate candidate genes, the regulatory elements of which could ensure constitutive and robust transgene expression in *S. mansoni*. Based on transcriptomics data (6), we became attentive to Smp_335990, a gene that appears to be strongly, ubiquitously (8), and stage-independently expressed (Supplementary Figure S1). Annotation in WormBase ParaSite (37) identifies Smp_335990 as a ubiquitin-like domain-containing protein (A0A5K4F9I1; UniProt). Protein structure prediction by Phyre2 (38) supported the annotation and identification as orthologous to UBB, human ubiquitin B (39–41) (Supplementary Figure S2). A 2,056 bp region upstream of the coding sequence of Sm*ubi* was scanned for the presence of hallmark promoter motifs (Figure 1A). Archetypal promoter and enhancer elements were identified (Supplementary Table S1). Promoter analysis using |the Neural Network Promoter Prediction tool (42) predicted two core promoters: at position 1,981 to 2,030 bp, promoter rank 1 (Supplementary Tables S1, S2) and 156 to 205 bp, promoter rank 2 (Supplementary Table S3). Next, these regions were analyzed in more detail with AliBaba2.1 (43), TRANSFAC 4.0 (44), YAPP eukaryotic core promoter predictor (45–48), and by comparison with known consensus promoter elements (49–51) (Supplementary Tables S2, S3). In the core promoter rank 1, typical sequence elements required for transcription initiation were present including TATA– and CAAT-boxes, the B recognition element (BRE, the transcription initiator element (INR), and the downstream promoter element (DPE). Consensus sequences of enhancer elements including Fushi tarazu (FTz and CAAT/enhancer-binding sites (C/EBPalp) were present as were binding sites for the serum response factor (SRF), hepatocyte nuclear factor 1 binding element (HNF-1), heat shock transcription factor 1 (HSF), and octamer-binding transcription factor 1 (Oct-1). The start-codon containing Kozak sequence, necessary or ribosome binding and initiation of translation (52) was identified *in silico* by comparing the potential Kozak sequences of *S. mansoni* genes known to be abundantly transcribed and sharing comparable transcription strengths (53, 54) and expression patterns (6, 54) to that of Sm*ubi*. For these genes, we predicted the consensus sequence 5’-NNT/CA/GNA/G/TATGNC/AN-3’ 6 bp downstream and 5 bp upstream of the translation start (Supplementary Table S2).

**Figure 1.**
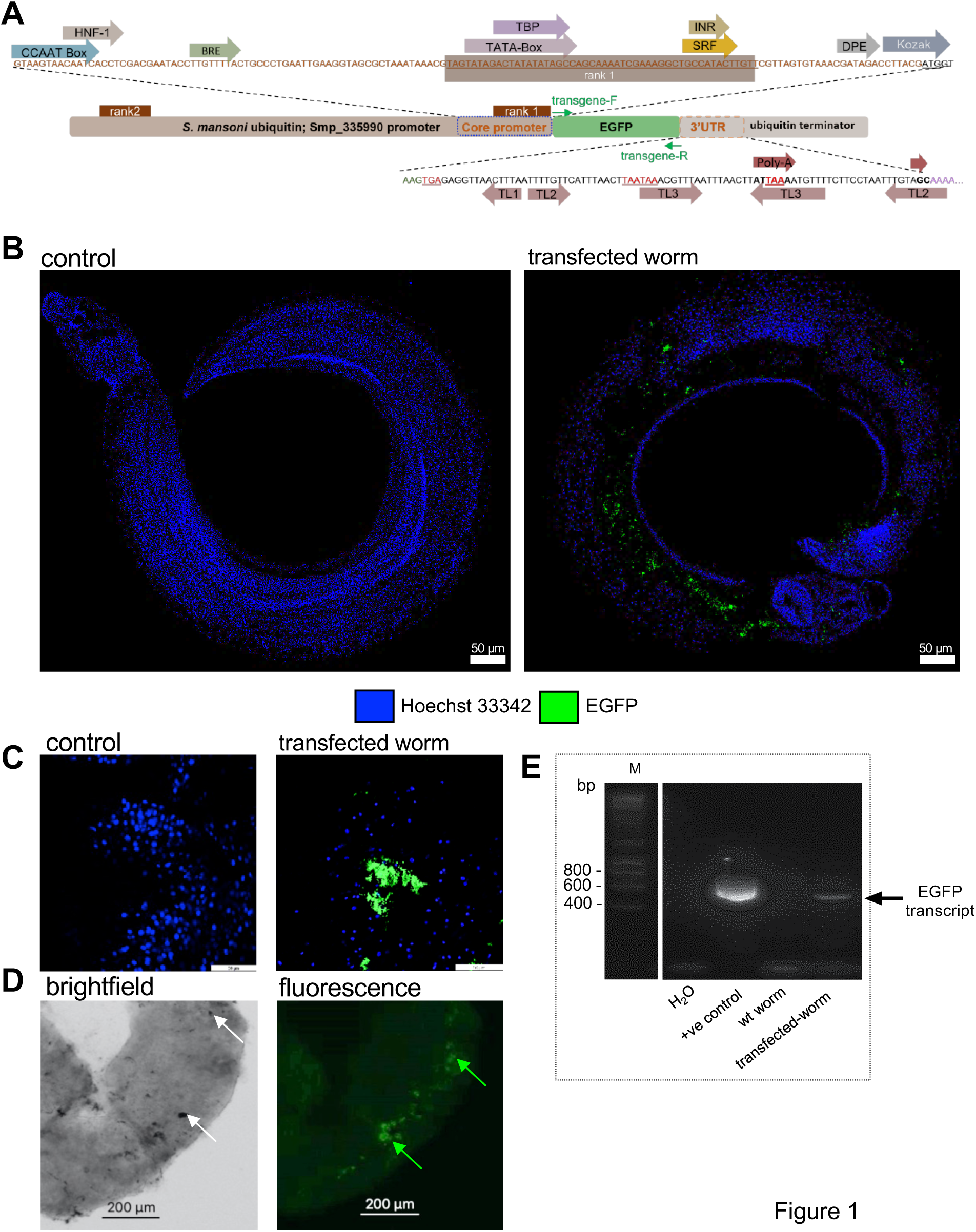
Bioinformatic prediction and experimental verification of the regulatory activity of *S. mansoni* ubiquitin (Sm*ubi*) promoter and terminator. **(A)** Overview of Sm*ubi* (Smp_335990) promotor and terminator elements (see Supplementary Tables S1, S2). The elements were identified using Berkeley *Drosophila* Genome Project Neural Network PromoterPrediction Tool, AliBaba2.1 TRANSFAC 4.0 (44), and YAPP eukaryotic core promoter predictor (45–48), respectively. Two core promoters were predicted (brown box and arrow) containing a CCAAT enhancer-binding site (light blue arrow), hepatocyte nuclear factor 1 binding element (NHF-1), B recognition element (BRE), TATA-Box-containing region (TBP), transcription initiator element (INR), serum response factor (SRF), and downstream promoter element (DPE). The presumptive terminator region included three terminator stem-loop structures and the polyadenylation signal, as predicted with the RNAfold Webserver Vienna tools (56) and the DNAFSMiner-DNA Functional Site Miner polyadenylation side prediction tool (55), respectively. Sm*ubi* promoter-induced EGFP reporter-gene expression was monitored in adult male *S. mansoni* worms, which had been transiently transfected by particle bombardment (PB) (86) with the pJC53.2_SmUbi-EGFP-SmUbi construct. Following cloning of the SmUbi-EGFP-SmUbi reporter-gene construct (using the rank 1 core promoter sequence) into the plasmid vector pJC53.2 (122), adult male schistosomes were transfected by PB. After 48 h, transfected and control worms were counterstained with Hoechst33342 (blue) and examined using CLSM **(Panels (B) and (C))**. Green fluorescing regions were seen while EGFP signals were not seen in control worms. Brightfield microscopy confirmed the presence of clusters of gold particles in the tissue (white arrows, **Panel (D)**). EGFP signals colocalized with the gold particles (green arrows, **Panel (D)**). **(E)** RT-PCR confirmed the presence of EGFP transcripts in transfected worms. No EGFP transcripts occurred in wild type (WT) worms (control) or in a control without template. +ve = positive RT-PCR control; H_2_O, control RT-PCR without template. Abbreviations: CLSM, confocal laser scanning microscopy; EGFP, enhanced green fluorescent protein.

We surveyed the potential terminator sequences including the 3’UTR (3, 5) of Sm*ubi* (Figure 1A; Supplementary Table S3). Applying the DNAFSMiner prediction tool for the analysis of functional sites in DNA sequences (55), a polyadenylation signal within the Sm*ubi* 3’UTR was apparent. RNA secondary structure prediction by RNAfold (56) revealed several stemloop structures that are typical for transcription termination (57).

### Schistosome ubiquitin promoter drives episomal EGFP

Next, to empirically test promoter activity of SmUbi-EGFP-SmUbi, adult male schistosomes were transfected with the plasmid construct by particle bombardment (PB) (Figures 1B, 1C and 1D), a ballistic gene-transfer method suitable for the transient transformation of adult and larval schistosomes (58, 59). Green fluorescing regions were apparent by 48 hours (Figure 1B, C, D) and EGFP transcription in these worms confirmed by RT-PCR (DNase I-treated RNA). (Figure 1E).

### Programmed KO at GSH1 in the schistosome egg

The *S. mansoni* GSH1 is AT rich (66%), which potentially constrains the value of the GC-content dependent Cas9 for programmed genome editing at this location. Cas12a is better suited than Cas9 for editing AT-rich regions. Furthermore, we showed previously that overlapping of three gRNAs combined with Cas9 cleaved the target DNA with resultant staggered (overhanging rather than blunt ended) strands (3, 21, 26). This outcome in turn led to marked increase of ∼70% of donor transgene integration and chromosomal repair by HDR in the schistosome egg. Accordingly, we hypothesized that DNA cleavage by Cas12a combined with a single crRNA, with releases staggered strands at cleavage, should enhance the efficiency of programmed gene editing. To address the hypothesis, we designed gRNAs for Cas12a and for Cas9 to cleave at the same target position (Figure 2A). These gRNAs also shared a similar predicted CRISPR/Cas-efficiency and absence of predicted off-target activity.

**Figure 2.**
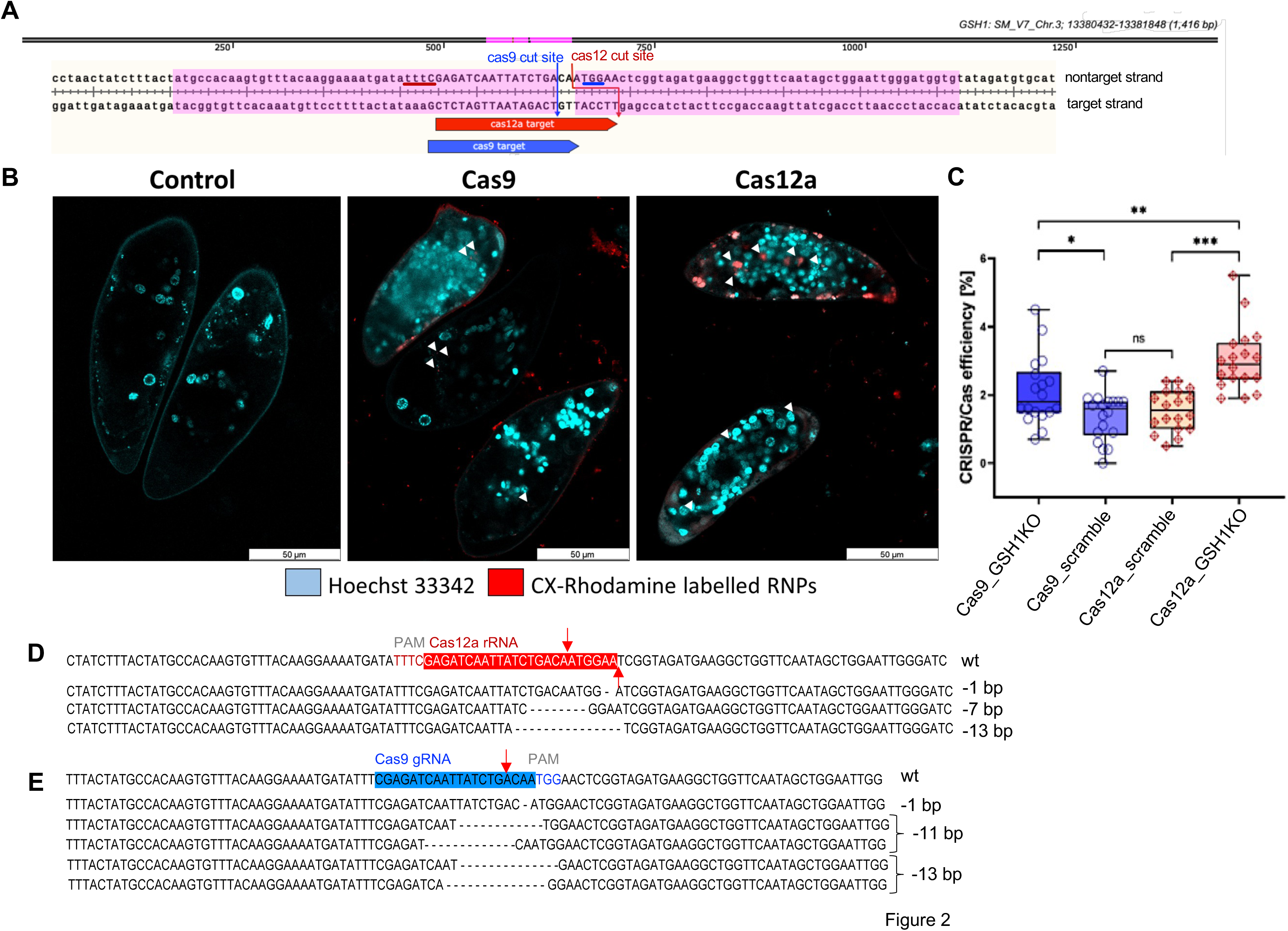
Comparison of CRISPR programmed gene knockout at genome safe harbor 1 (GSH1) in the egg stage *Schistosoma mansoni* by Cas12a versus Cas9. **(A)** Schematic overview of the GSH1 (SM_V7_Chr3; 13380432-13381848; 1,416 bp) target sites for Cas9 (blue arrow) and Cas12a (red arrow). Cas9 cleavage releases blunt ended DNA strands whereas Cas12a releases overhanging strands. Cleavage at GSH1 was predicted on the non-target DNA at the position illustrated. GSH1 regions highlighted in magenta represent the short homology arms, 50 nt, of the donor template. **(B)** LEs were subjected to electroporation in the presence of CX rhodamine labeled RNPs (red), after which they were counterstained with Hoechst33342 (blue) and examined 24 hours later by CLSM. A majority of Cas9 RNPs accumulated just below the eggshell, a dissimilar pattern to the Cas12a RNP-treated LEs, where signals were also evident in the deeper tissues of the egg. **(C)** Editing efficiency (InDel abundance) of Cas9 and Cas12a RNPs at GSH1 as assessed by TIDE (62). The editing efficiencies of GSH1-targeting RNPs normalized to the wild-type sequence (KO-Cas9, dark blue bar; KO-Cas12a, red bar) and RNPs non-specific to the *S. mansoni* genome (scramble-Cas9, purple bar; scramble-Cas12a, orange bar), respectively, are shown. The editing efficiency of LEs transfected with RNPs was significantly higher to transfection with scramble control PNPs (*P < 0.05, **P < 0.01, ***P < 0.001; *t* test, *n* = 9). Compared with Cas9-RNPs, Cas12a-RNPs (red) induced significantly more efficient programmed KO. **(D)**, Representative NGS reads for Cas12a and Cas9, respectively, of alleles with frequently seen indel profiles at the programmed cleavage position (red arrows) (n = 3). Abbreviations: CLSM, confocal laser scanning microscopy; KO, knock-out, ns, not significant; wt, wild type.

To establish genome editing in schistosomes by chromosomally integrated reporter genes, a suitable life stage(s) of the parasite should be targeted, a compatible method to introduce gRNAs, replacement constructs in case of KI experiments, and the optimal enzyme to efficiently edit the target region. To this end, the egg stage was used, given its suitability for electroporation-mediated transformation. First, the accessibility of LEs by RNPs of both enzymes was determined using an RNP-tracking approach, where fluorescence-labelled RNPs were introduced into LEs by electroporation. Each guide RNA – sgRNA of Cas9, crRNA of Cas12a) was labeled with CX-rhodamine before to assembly of RNP complexes with *Streptococcus pyogenes* Cas9 (SpCas9) and *Acidaminococcus* sp. Cas12a (AsCas12a) nucleases (26, 60, 61). LE were transfected with the RNPs by square-wave electroporation. Confocal laser scanning microscopy (CLSM) was used to monitor entry of RNPs into the eggs; by 24 hours, entry of the RNPs of Cas9 and of Cas12a was confirmed in the larval tissues of the schistosome egg, which colocalized with Hoechst 33342-stained nuclei (Figure 2B). With Cas12a, RNPs had penetrated the eggshell and were detected in or near larval cells. With Cas9-RNP, while the majority of Cas9-RNP traversed the eggshell as with Cas12a RNPs, comparatively less labeled Cas9 RNP was evident in larval cells. Based on these micrographs, we conclude that electroporation propelled both RNPs formed into the egg, but that Cas9 RNPs accumulated proximal to the eggshell whereas Cas12 RNPs were more widely distributed through the developing larva within the eggshell.

Next, after establishing that RNPs of both Cas nucleases entered the schistosome egg, LEs were transfected with unlabeled RNPs to examine the accessibility of GSH1 and the performance these two RNP types. The eggs were collected 10 days after transfection, gDNA extracted, and PCRs performed using the gDNAs as template. TIDE analysis was undertaken on Sanger sequence reads (26, 62), of nine independent biological replicates (replicates 1-9). TIDE revealed programmed KO efficiency of 2.2±0.9% and 3.0±0.9% for Cas9-RNP and Cas12a-RNP, respectively (Figure 2C). The wild type (untreated), the electroporation control (mock), and LEs treated with RNPs assembled with a scrambled sequence crRNA, GCACTACCAGAGCTAACTCA (63) were included to assess natural sequence variation and CRISPR/Cas off-target activity at GSH1. To normalize for the background artefacts of the sequencing procedure, we subtracted the editing efficiency of the experimental groups (RNP treatment) with the values for wild type and mock controls (∼0.1%). The mutation rate in LEs treated with RNPs with the scrambled (irrelevant) crRNA was 1.4±0.7%, significantly lower than Cas9 (*P* < 0.05) and Cas12a (*P* <0.001) programmed to cleave GSH1 (Figure 2C).

Next, we combined gDNAs (after TIDE analysis) from sample numbers 1-3, 4-6 and 8-9 for target amplicon NGS sequencing. NGS paired-reads (∼100,000 reads for each sample) from these three independent NGS libraries were analyzed using CRISPRessV2 and using Rgen Tools confirmed the precise, on-target editing catalyzed by both enzymes (Figure 2D and 2E) (64–66). Nucleotide deletion assessment on expected cleavage site showed alleles with deletions of 1, 7 and 13 nt in length by Cas12a, and 1, 11 and 13 nt by Cas9 (Figure 2D, 2E). These allele genotypes were similar in all three NGS libraries.

A similar programmed KO outcome was seen for Cas12a at GSH1 in adult stage paired female and male schistosomes, although here significantly higher editing efficiency was observed with Cas12a compared to Cas9-treated worms (*P* < 0.01; Supplementary Figure S3A). The size of deleted DNA fragments of GSH1 ranged from 9-18 nt in the Cas12a-treated group, with 1 nt deleted by Cas9 (Supplementary Figure S3B, S3C). In review, these findings confirmed programmed KO with both Cas nucleases in *S. mansoni* eggs and in adults, but they convincingly confirmed significantly higher editing efficiency of Cas12a over Cas9 at GSH1.

### Programmed KI using a transgene flanked by microhomology arms as the repair template

We investigated whether Cas9 and Cas12a equally efficiently insert the transgene into GSH1 at programmed CRISPR catalyzed HDR. Given that 5’ C6 amine-PEG10(C6-PEG10)-modified donors (with ∼50 nt HA) enhance HDR efficiency in HEK293 cells (60, 61), we likewise tested chemically modified double-stranded DNA donor as the template for HDR insertion into GSH1, specifically amine C6 with PEGylation (PEG 10 kDa) at 5’ termini of the donor bearing minimal length (micro)homology arms each of 50 nt in length. PCR amplicons of the SmUbi-EGFP-SmUbi transgene were employed as the donor template. RNPs assembled with Cas9 and Cas12a were used and compared in this assay. For both nucleases, transgene integration at GSH1 was achieved at ratios of 1:1:1 (w:w:w) of enzyme: Cas9 sgRNA or Cas12a crRNA: donor EGFP in each case. The 5’ HA of the donor template contained the Cas12a PAM recognition site (TTTC) and a part of the Cas12a target sequence, one base of the Cas12a PAM TTTC was altered to T<u>C</u>TC to eliminate the Cas12a specific PAM (Figure 3A). This was undertaken to impede donor DNA cleavage by Cas12a. In addition, the design of the gRNAs and selection of the cognate PAM recognition site of both Cas9 and Cas12a were aimed at ensuring programmed KI by using the identical microbiology arms, targeting GSH1, on the donor DNA amplicons. This experimental design was achieved by selecting predicted and near-identical cleavage sites of the nucleases at GSH1 (Figure 3A).

**Figure 3.**
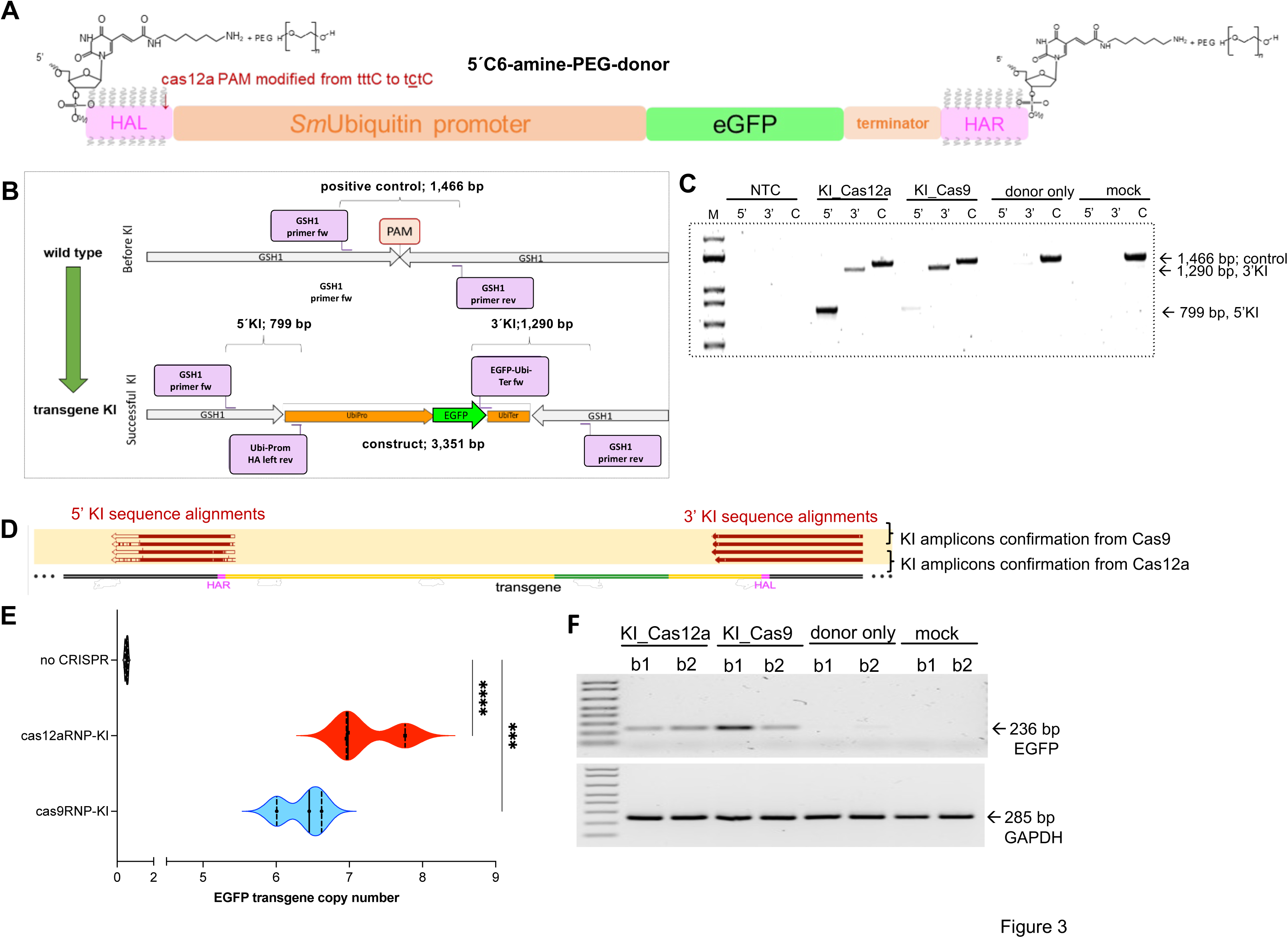
RNP-mediated chromosomal integration of the SmUbi-EGFP-SmUbi transgene. **(A)** Schematic representation of the transgene construct: the *S. mansoni* ubiquitin promoter (SmUbi, 2,056 bp) and terminator (Smubi, 302 bp) shown in orange flank the EGFP reporter (green; 717 bp). The terminal microhomology arms – HAL and HAR, 50 nt each in length, are indicated in pink. Chemical (5’-C6 amine-PEG10) modifications to protect the transgene from cellular degradation ((60) are indicated. Last, to avoid Cas12a-mediated cleavage of the SmUbi-EGFP-SmUbi donor template, the 5’-TTTV-3’ Cas12a PAM within the microhomology arms was altered to 5’-TCTV’3’ (red letters). **(B)** Schematic depiction of the GSH1 locus before and after KI of the SmUbi-EGFP-SmUbi (orange and green) reporter construct, with the position of the PAM indicated (red) and showing the primer binding sites for the KI analyses. **(C)** The success of the chromosomal integration of the SmUbi-EGFP-SmUbi donor template was demonstrated by PCR amplicons that span the 3’– and 5’-integration sites. The wild type GSH1 locus served as a control, whereas the complete transgene integration region was not expected to be amplified due to limitations of the PCR. DNA fragments specific for the 5’and 3’ integrations of the donor template were obtained by PCR and amplicons sequence reads obtained with by Sanger direct sequencing. **(D)** For both Cas9 and Cas12a, alignment analyses of reads of these integration region amplicons confirmed the integration of the transgene (yellow and green) into GSH1 (grey), as indicated by red bars. **(E)** Estimation of EGFP transgene copy number in gDNA (at day 10 after transfection) obtained by interpolation from the external standard curve plotted for plasmid DNA, as described (26). Transgene copy numbers in the Cas9 and Cas12a groups were significantly higher than LEs transfected with the donor template only (no RNP). KI analysis of the Cas9 group showed a broad range of EGFP copy numbers compared to the Cas12a group (2Way ANOVA, \*\*\**P* = 0.0001, **** *P*< 0.0001). Significantly fewer copies of residual (episomal) donor DNA were present in the donor-only control; grey violin plot). **(F)** RNA/cDNA of eggs of were analyzed by RT-PCR to monitor EGFP transcription, which demonstrated EGFP transcripts (236 bp) in LEs of the KI treatment groups using Cas12a and Cas9. DNase I treated-RNA served as a control for the RT-PCR to eliminate the contaminated DNA and/or remaining donor. Lanes b1 and b2 indicate the outcome from two independent biological replicates. EGFP transcripts were not detected in the controls (top panel). The SmGAPDH gene GAPDH (Smp_0569701, 285 bp, bottom panel) served as the reference control for cDNA synthesis and RT-PCR.

We co-electroporated RNPs and this donor DNA template into LEs of *S. mansoni*. Mock electroporated wildtype LEs served as a transfection control; LEs transfected with the donor template in absence of the RNP. Ten days after transfection, the eggs were collected, gDNA isolated, and investigated for transgene integration. We designed two primer pairs that, firstly, amplify the 5’-integration site (5’-KI) including sequences within the GSH1, i.e., the SmUbi-promoter region of the transgene (799 bp, Figure 3B) and, and secondly, 3’-KI including the 3’-portion of EGFP and GSH1 (1,290 bp, Figure 3B). Using this strategy, we amplified 5’-KI and 3’-KI integration fragments and thereby demonstrated GSH1 editing and KI by both nucleases (Figure 3C). Amplicon band intensity of 5’-KI amplicons in the Cas9 group was less intense than for Cas12a-treated LE (Figure 3C).

Nonetheless, alignment analyses using Sanger sequence reads, obtained from the purified 5’-KI and 3’-KI amplicons, confirmed programmed transgene KI into GSH1 (Figure 3D). As expected, 5’-KI and 3’-KI fragments were not amplified using DNA from the control groups. A 1,466 bp fragment of the non-edited GSH1, amplified in all experimental groups, served as a PCR control. Next, we quantified the copy number of SmUbi-EGFP-SmUbi transgenes integrated into the genome in edited LEs compared to LEs, which were transfected only with the donor template in the absence of RNPs (Figure 3E). For this, the external standard curve design was employed to estimate integrated-EGFP in gDNA of the schistosome eggs (n = 3) as described (26). The standard curve is based on linear regression (y = 3.3857x+35.093, R^2^ = 0.9979) established by a logarithm (20)-transformed initial DNA input, the pJCR53.2-SmUbi-EGFP-SmUbi plasmid was used as the dependent and the Ct value from the qPCRs as the independent variation (not shown). We estimated the EGFP transgene copy number in GSH1 by converting the observed Ct values to the logarithmic copy numbers using the equations obtained from the standard curve. There were 6.594±0.196 and 7.235±0.458 EGFP copies in ∼10 ng DNA resulting from Cas9 and Cas12a, respectively (*P* < 0.01). The control group (donor template only) showed 0.516±0.082 copies (10 ng of gDNA), which probably reflected residual non-integrated SmUbi-EGFP-SmUbi donor DNA. Together with this finding, EGFP transcripts were not detected by RT-PCR of the RNA/cDNA of the control groups, and only found in KI in DNase I-treated RNA samples (Figure 3F). In summary, the copy number of integrated SmUbi-EGFP-SmUbi was significantly higher with Cas12a than Cas9 catalyzed KI.

### EGFP expression in the miracidium

CLSM was employed to evaluate reporter gene expression in eggs/miracidia following programmed KI at days 5 and 10 after transfection. Emission spectra were analyzed as described (21), which revealed EGFP fluorescence in cells of eggs that contained a miracidium. Fluorescence intensity of eggs emitted at 509 nm was quantified, which revealed significantly more EGFP-positive eggs in the Cas12a group than the Cas9 group at days 5 and 10 after electroporation (Figure 4A). For this analysis, eggs were scored as EGFP-positive if they included a miracidium with >50% EGFP expression in the investigated section plane (two dimensions). From six biological replicates, the percentages of EGFP-positive eggs were 43.6 ±19.9% and 26.3±7.2% of eggs were EGFP-positive in the Cas12a group and 20.2±10.1% and 21±16.2% of eggs of the Cas9 group were EGFP-positive at days 5 and 10 (*P* < 0.01 to 0.001) (Figure 4B). Differences in growth and survival were not noted between transfected eggs and eggs from the control groups. Viability of treated eggs was confirmed by hatching of miracidia (not shown).

**Figure 4.**
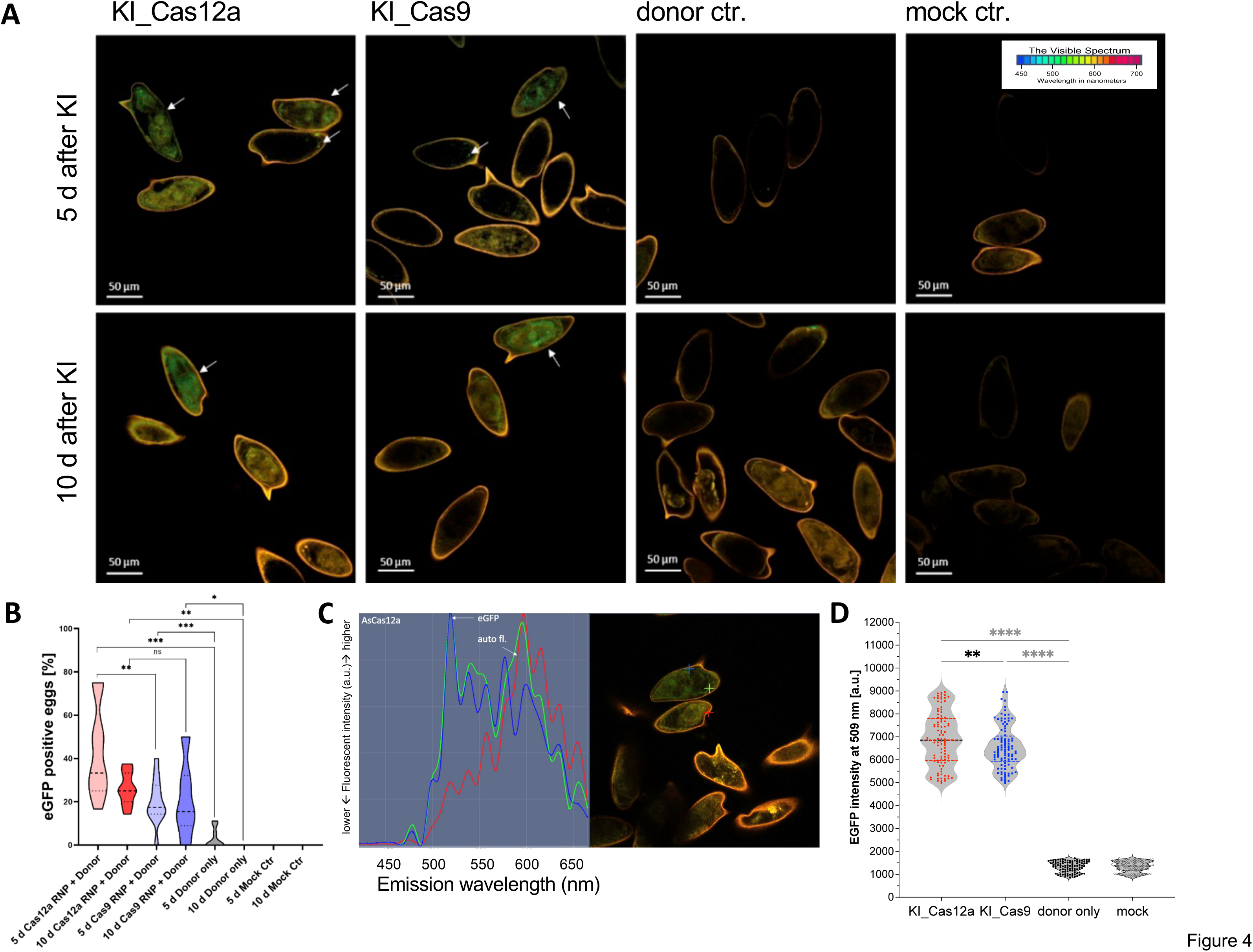
CRISPR/Cas12a more efficiently catalyzed transgene KI than Cas9. **(A)** CLSM micrographs of miracidia with green fluorescence within the eggshell at days 5 and 10 days. Orange color indicates autofluorescence. Scale bar: 50 µm. **(B)** The proportion of EGFP-positive eggs in the groups was assessed by CLSM at 5 and 10 days: ∼10,000 eggs of each biological replicate (*n* = 6). Cas12a (red) displayed higher EGFP-positive rates than Cas9 (blue) after both days 5 and 10. Both Cas12a– and Cas9 transfected eggs contained significantly more fluorescing larvae (miracidia) than the donor only control group (grey). EGFP fluorescence was not seen in control, untreated eggs (\**P*<0.05, \*\**P*<0.01, \*\*\**P*<0.001; *t* test). **Panels (C) and (D)** Verification of EGFP signals in LEs transfected with Cas12a RNPs after 5 days. Using the Zeiss Zen blue 3.4 software, different positions of the fluorescence spectrum were sampled to distinguish EGFP fluorescence from the background autofluorescence (orange). The typical EGFP spectrum (left part; blue and green lines) was evident in the larvae (right panel, green and blue + signs) and was distinct from the autofluorescence spectrum (right panel, red + sign) of the egg, which mainly localized in the eggshell (red + sign). The EGFP-positive miracidium from donor only and mock treatment groups showed fluorescent intensities of ∼1333.84±235.38 a.u. and 1331.12±1169.35 a.u., respectively. The EGFP-KI mediated by Cas12a exhibited higher signal intensity than Cas9: 6897.25±1169.31 a.u. and 6548.71±941.11 a.u., respectively (one-way ANOVA \*\*\*\**P*< 0.0001, **P = 0.0079, *n* = 100). The spectral intensity of the autofluorescence in donor only (without RNPs) and mock electroporated eggs were not significant different. Abbreviations: auto fl., auto fluorescence; CLSM, confocal laser scanning microscopy; CTRL, control; HA, homology arms; KI, knock-in; LE, liver eggs; primer fw, forward primer; primer rev, reverse primer.

A weak EGFP signal at 509 nm was detected in LEs transfected only with the donor only (Figure 4B). Subsequent RT-PCR analyses with egg RNA/cDNA of all groups confirmed EGFP transcripts in the treatment groups but not in the control groups, except a weak EGFP background signal in LEs transfected with the donor template in the absence of RNPs (Figure 3C).

The EGFP intensity at 509 nm, the peak emission wavelength for this reporter in miracidia was determined at day 10 days after electroporation. One hundred eggs in each treatment group were investigated for EGFP signal-intensity of the viable miracidium within the eggs, using a formula that subtracted autofluorescence signals (21). The EGFP-specific signal intensities from the eggs, which resulted from SmUbi-EGFP-SmUbi integration, were significantly different between the CRISPR-Cas9 and –Cas12 groups with mean ± SD intensities of 6548.71±941.11 a.u. and 6,897.25±1,169.31 a.u., respectively (Ordinary one way ANOVA, P value 0.0079). EGFP signals were not detected in the control groups; donor only, and mock transfections; these groups exhibited minimal fluorescence at 1333.84±235.38 a.u. and 1331.12±232.35 a.u., respectively (Figure 4D).

## DISCUSSION

Neglected tropical diseases afflict more than one billion people, frequently residents of in some of the world’s most impoverished communities and regions. Despite the devastating health consequences, these diseases receive too little attention from global funding agencies, pharmaceutical companies and biotech research and development. They can be challenging to study given many are not laboratory-accessible, they can have multiple developmental hosts in the life cycle, they may hypoxic or other non-standard physicochemical culture constraints, and they possess complex diploid genomes (67–70). These impediments, notwithstanding, to continue to advance functional genomics of helminths (71–73), here we report progress in genome editing at the genome safe harbor site GSH1 in *S. mansoni* (21) by comparing the editing capacities of Cas9 and Cas12a, and optimized RNP delivery conditions and donor transgene structure. We observed efficient integration of a reporter gene into GSH1 using 5’-C6-PEG10 modified 50 bp microhomology arms by programmed CRISPR-Cas9/Cas12a-induced HDR repair, as confirmed by amplicon sequencing as well as fluorescence reporter gene activity.

Knockout models are widely used for functional studies of gene(s) of interest. However, in the *omics* era of schistosome and other helminth research, facile protocols for stable transformation are not readily available. CRISPR/Cas gene editing is now the state of the art for genome manipulation (74), and there are reports involving CRISPR gene editing in trematodes (26, 71, 75, 76). Genome editing was limited to eggs, and genome manipulation probably accessed the outer cell layers during embryogenesis (36, 77, 78). Yet, to establish stably transformed lines, approaches need optimize germline transgenesis. We had established a protocol for RNA-programmed Cas9-catalysed editing at GSH1 in *S. mansoni* that used multiple, overlapping guide RNAs, after computational prediction of the genome coordinates for this GSH based on genome annotation, chromosome structure, and experimental validation (21). Here, we boosted both this method and enhanced gene-editing efficiency using a single gRNA-programmed Cas12a in the presence of a donor repair template bearing chemically modified, microhomology arms, which, in combination, enhanced programmed KI of the transgene.

Both egg and adult stages of *S. mansoni* were transfected by electroporation of Cas-containing RNPs. Treatment of adult schistosomes with RNPs transferred by electroporation failed to achieve a penetration and distribution RNPs in all tissues (75). Indeed, exposure of adult stage schistosomes to CRISPR by electroporation can be expected to result in genetic mosaicism. Our findings conformed with earlier reports of deletion mutations at CRISPR targets in LE, schistosomula, and adult schistosomes (26, 71, 75, 79, 80). Nonetheless, the outcome can be informative for proof-of-principle of genome editing or as an alternative for RNA interference of genes transcribed in the (sub)tegument.

To overcome these problems and to establish a genetically homogeneously modified multicellular parasite, it is necessary to target life stages with a limited number of cells, or a stage rich in stem/germinal cells such as the schistosomulum (9), or at an early developmental stage such as the zygote (36). For this purpose, we aimed to introduce RNPs into LEs, isolated from livers of infected rodents, and which consisted of egg subpopulations representing a spectrum of developmental stages. Notably, this LE population includes non-embryonated eggs, consisting of the zygote and a few vitelline cells (36). Confocal microscopy of LEs transfected with RNPs, which incorporated rhodamine-labelled Cas9-gRNAs (75) or Cas12a-gRNAs, demonstrated the suitability of this approach to permeate the eggshell, and where Cas12a RNPs showed a higher efficiency to access embryonic cells. Moreover, transfection of the zygotes at an early developmental stage was visualized by RNP-tracking experiments (Supplemental Figure S5). The higher efficiency might reasonably be explained by the size difference of RNPs given that the Cas12a-RNP complex (Cas12a: 150.9 kDa) was smaller than Cas9-RNP complex (Cas9: 161.3 kDa) (30), likely enabling a comparatively better uptake of through the pores in the egg (81). Moreover, Cas12a gRNA at 41-44 nt is shorter than the Cas9 gRNA (100 nt) (30). RNPs of both nucleases were detected in numerous cells of the eggs but a limitation of this approach was that not every cell appeared positive for the RNPs. To optimize this, alternative transfection methods such as lipid nanoparticles (82) and other methods (59, 83–87) could be tested. Alternatively, one could envisage targeting other larval stages such as the sporocysts to enhance transfection efficiency (75).

In addition to these differences in the efficiency of eggshell penetration, analysis of the deletion frequencies of GSH1 revealed a significantly higher editing efficiency of Cas12a over Cas9 in eggs and adult schistosomes. Indeed, the editing efficiency of Cas12a with a single gRNA was comparable to the efficiency of Cas9 with two to three gRNAs (21). The differences might result from i) the efficiency bias of Cas12a for penetration of the eggshell, ii) differences in the cleavage profiles of the nucleases (Cas9, blunt ends; Cas12a, overhanging ends) (30), iii) divergent efficiency of NHEJ, which relies on the gRNA sequence and cell-cycle stage of the cells (88–90), and/or iv) combinations of these factors.

Intriguingly, the deletion patterns of Cas12a and Cas9 differed between the egg and adult suggesting that DNA repair preferences differ during development in this schistosome. The egg in utero is comprised mostly of S4 vitelline cells (91), the fertilized oocyte in the early phase, and the miracidium with surrounding acellular and cellular layers – Reynolds’ layer and von Lichtenberg’s envelope, respectively (92) at a late stage (36, 93). We speculate that DNA repair profiles differs among these cell types and developmental stages. Findings in other metazoan species support the conjecture (94–97). Aiming to buttress support for this hypothesis, we performed a first i*n silico* approach to identify orthologs in *S. mansoni* of genes participating in double stranded DNA break repair, NHEJ and HDR, in *S. mansoni* based on data available for the model nematodes, *Caenorhabditis elegans* and *Pristionchus pacificus* (Supplementary Table S5, Supplementary Figure S5) (98–103). (Notably, schistosome proteins predicted to participate double-stranded break (DSB) repair did not exhibit marked identity to mammalian DSB-repair pathway members (90, 104).) Expression patterns of *S. mansoni* protein-coding genes, predicted to be involved in DSB repair (7, 53, 54) showed clear differences from the tissue-preferential transcript profiles of genes attributable to NHEJ (e.g., SmCKU70, SmCKU80) or HDR (e.g., SmHIM6, SmRAD51, SmRPA). NHEJ-associated genes were transcribed throughout the schistosome tissues whereas HDR-associated proteins were preferentially transcribed in the gonads (7, 97, 105). Genes predicted to be involved in HDR, including SmRAD51, SmRPA and SmBRC1, were not transcribed in S4 vitellocytes (54). These variations likely underlie the divergent mutation rates and alleles between the egg and adult stages. In addition, HDR is the prominent repair pathway expected to operate in proliferating cells including embryonic cells (90), and where indels are not expected to accompany perfect DSB repair (95). Correspondingly, increased transcript levels of HDR-associated genes might be anticipated in the parasitic stages of the developmental cycle including the sporocyst, which undergo intense cellular proliferation (53, 106). By contrast, terminally differentiated cells such as the S4 vitelline cell likely rely on NHEJ (90, 104) for DNA repair (88, 94).

We also confirmed the utility of microhomology (50 nt) arms for CRISPR-Cas-induced HDR repair, as demonstrated by constitutive reporter gene expression in the embryonating miracidium. Cas9 and Cas12a were each introduced into NLE the presence of a 5’C6-PEG10-modified donor template coding for EGFP under control of the endogenous ubiquitin promoter of *S. mansoni* (21). We analyzed this promoter in detail and identified several core promoter elements (49, 51, 107–109), including a schistosomal Kozak sequence (52). The promoter elements may contribute to the constitutive transcriptional activity of the cognate ubiquitin gene in the developmental cycle (53). Constraints on transgene cargo size delivered by HDR are well known (21, 110, 111). However, the use of minimal length homology arms of 50 bp in length flanking the transgene, i.e., *microhomology* arms as deployed here, can facilitate insertion of transgene cargo up to 9 kb in length (61, 112–114).

For reporter gene integration, we used a chemically modified (5’-C6-PEG10) donor DNA template, modifications aimed at preventing DNA degeneration by endogenous nucleases and also known to enhance HDR in human cell lines (HCT116, HEK293T, H1, and WTC G3) (61). Superior KI performance has been reported for the donor template in some human cell lines when the template combined 5’C6-PEG10 modification and short HAs (60, 61). Transgene insertion into GSH1 using this adapted approach for schistosomes represents a significant optimization over our original protocol (21) as both HAs were substantially shorter and dsDNA could be used instead of a single-stranded DNA donor. Likewise, with an RNP assembled with Cas12a and a single crRNA, we achieved comparable and higher integration rates with the modified donor template compared with Cas9 and up to three gRNAs (21). Notably, Cas12a looks to be well-suited for *S. mansoni*, given its AT-rich genome (3), and given that Cas12a has a T-rich specific PAM, 5’-TTTN-3’ (30). In overview, Cas12a exhibits a divergent flexibility to Cas9, which is constrained in its target sequence range in the schistosome genome due to its CG-rich PAM, 5’-NGG-3’.

Optimized genome editing for schistosomes can be expected to hasten research progress in functional genomics of these helminths. Our recent report defining a panel of genome safe harbors in *S. mansoni* provided a salient innovation for CRISPR-based functional genomics and transgenesis in *S. mansoni* (21). Now, we have also overcome other earlier limitations through deployment of Cas12a as the RNA-guided nuclease and the modification of the donor transgene with chemically protected, diminutive length (micro)homology arms. Along with loss-of-function mutation, these improvements will allow gain-of-function approaches by, for example, integrating selectable marker genes into a GSH as an alternative for, or in addition to, fluorescence reporters. This would enable maintenance of transgenic lines with drug selection pressure. Indeed, oxamniquine showed selectivity for sporocysts and adult schistosomes (75). Beyond that, this model should be useful during development of new interventions for schistosomiasis and other similar neglected tropical diseases. This method also opens the way for studying genes inaccessible to RNA interference (17, 19, 115). Last, we posit that this new workflow provides a blueprint for the design of advanced, tissue-stage investigation and contribute to deciphering the complex biology of schistosomes, which in turn, provides a role model for platyhelminths generally.

## MATERIAL AND METHODS

### Maintenance and isolation of *S. mansoni* developmental stages

Mice (female, Swiss Webster) exposed to ∼180 cercariae of *S. mansoni* were provided by the Schistosomiasis Resource Center, Biomedical Research Institute, Rockville, MD. The mice were housed at the Animal Research Facility of George Washington University (GWU), which is accredited by the American Association for Accreditation of Laboratory Animal Care (AAALAC no. 000347) and has the Animal Welfare Assurance on file with the National Institutes of Health, Office of Laboratory Animal Welfare, OLAW assurance number A3205. All procedures employed were consistent with the Guide for the Care and Use of Laboratory Animals. The Institutional Animal Care and Use Committee at GWU approved the protocol used for mice in this study. Mice were euthanized at about day 49 following infection. Schistosomes were recovered by hepatic portal vein perfusion with 150 mM sodium chloride, 15 mM sodium citrate, pH 7. Thereafter, the worms were washed with 1× phosphate buffer saline (PBS), pH 7.4, 2% penicillin/streptomycin/fungizone (PSF) and maintained thereafter in DMEM, 10% heat inactivated fetal bovine serum (FBS) (Corning Life Sciences, Corning, NY) supplemented with 2% antibiotic/antimycotic (Corning) in the atmosphere of 5% CO_2_, 37°C incubator (Panasonic, Newark, NJ) (116).

Adult schistosomes were processed within 24 h following recovery from the mice. To prepare *S. mansoni* eggs, mouse livers were rinsed in 70% ethanol, then twice washed with 1× PBS, 2% PSF, and finally homogenized in 1× PBS, 2% PSF in a mechanical blender. The homogenate was incubated with clostridial collagenase (0.7 mg/ml) at 37°C for 18 h, after which cellular debris was removed using sieves of 250 μm and 125 μm mesh (Gilson, Middleton, WI). Sequentially, followed by sucrose/Percoll gradient centrifugation, eggs were recovered (117). The eggs, termed “liver eggs” (LE), were maintained at 37°C in 20% FBS/ 2% PSF/ DMEM, and electroporated with CRISPR/Cas materials with or without donor (below) within 24 h.

### *In silico* analyses of the ubiquitin regulatory promoter elements

Enhanced green fluorescent protein, EGFP (118), under the control of the promotor and terminator sequences of Smp_335990 (119), an abundantly transcribed ubiquitin gene of *S. mansoni* (Sm*ubi*) (Supplementary Table S1, Supplementary Figures S1, S2) was used as a reporter for transfection/transformation analyses. Nucleotide sequences of upstream and downstream untranslated regions (UTRs) of Smp_335990 were retrieved from WormBase ParaSite (37, 119), version 7 of the draft genome of *S. mansoni* (3, 5). To confirm this annotation, we used the protein homology/analogy recognition engine Phyre2 with the protein structure modelling mode, http://www.sbg.bio.ic.ac.uk/~phyre2/html/page.cgi?id=index (43). To predict the promoter region of Sm*ubi*, adjacent sequences up to the three kb upstream of the start codon were scanned. Promoter elements were identified by the Berkeley *Drosophila* Genome Project Neural Network Promoter Prediction Tool, using a minimum promoter score of 0.8 for eukaryotic promoters (44), https://www.fruitfly.org/seq_tools/promoter.html. In addition, we used the AliBaba2.1 TRANSFAC 4.0 transcription factor binding site prediction tool, http://gene-regulation.com/pub/programs/alibaba2/ (45) to identify potential transcription factor binding sites. We analyzed potential promoter sequences also by AliBaba2.1 TRANSFAC (46), applying the following parameters: the Pairsim function was set to 50, the matrix width set to 10 bp, a minimum number of overlapping predicted transcription factor binding sites was limited to a total of four for each site with a minimum sequence conversation of 75% coverage, and the factor class was set to 4.

In addition, the sequence was analyzed for canonical core promotor elements by the YAPP eukaryotic core promoter predictor using a cut-off score of 0.8. To conduct the promoter analysis, the sequence including the predicted core promoter fused with the sequence of EGFP (717 bp) was used to analyze potential transcription start sites (TSS). Subsequently, TSS prediction was performed by YAPP (48, 50). To identify the schistosomal Kozak consensus sequence, we performed a comparative analysis applying the AGAT software-package to determine also potential Kozak sequences of other genes in the *S. mansoni* genome version 9. To this end, ubiquitously and abundantly transcribed genes were screened for the presence of potential Kozak sequences and patterns like Sm*ubi* (54). Specifically, the sequences of Smp_009580, Smp_106930, Smp_042160, Smp_056970, Smp_054160, Smp_182890, Smp_099870, Smp_179300, Smp_111340, Smp_090120, Smp_072330, Smp_003770, Smp_017430, Smp_040130, Smp_155060 and Smp_335990 were compared (120).

In addition, the 3’UTR of Sm*ubi* defined by WormBase ParaSite (37, 119), version 7 of the draft genome of *S. mansoni* (3, 5), was analyzed. To identify the polyadenylation signal, 600 bp downstream from the stop codon were analyzed with the DNAFSMiner-DNA Functional Site Miner polyadenylation side prediction tool (55).

Subsequently, the RNA secondary structure of the predicted 3’UTR was calculated, including the stop codon and predicted rank 1 (highest) polyadenylation site within *in silico* added poly-adenosine tail (10 adenosines). The investigation of potential RNA secondary structures characteristic for translation terminator-loops was done using the RNAfold Webserver Vienna tool, which calculates secondary structures by applying the minimum free energy (MFE) model and activated partition function, while the model avoids RNA folding with isolated base pairs, http://rna.tbi.univie.ac.at (55, 121).

### Reporter gene

Based on pJC53.2 (plasmid no. 26536, Addgene repository) (122), we cloned the reporter gene construct consisting of EGFP and the promoter and terminator sequences of Sm*ubi* (Figures 1A to 1D; Supplementary Table S1). One µg pJC53.2 was digested with *Ahd* I (New England Biolabs (NEB), Ipswich, MA) in 50 µl 1× CutSmart buffer (NEB), 120 min at 37°C, DNA fragments released (two fragments) were separated by agarose gel electrophoresis, and the 3,240 bp fragment recovered from the gel (Monarch Gel Extraction Kit, NEB, Ipswich, MA). Components of the transgene construct, i.e., the full-length EGFP coding region, putative promoter (5’Ubi) and terminator (3’Ubi) of schistosome ubiquitin were amplified by PCR (primers in Supplementary Table S4). Primers for the amplification of the 5’Ubi and 3’Ubi were designed according to data base information of WormBase ParaSite V7 for Smp_335990 (37, 119). Amplicons were synthesized in 20 µl reaction volume with either 100 ng gDNA or 10 ng pET-30a_SmDHFR-EGFP (the EGFP-DNA fragment kindly gifted by Dr. Jim Collins) as template for the reporter gene EGFP (717 bp). Expected sizes of the fragments for 5’Ubi and 3’Ubi were 2,056 bp and 580 bp, respectively. PCRs were undertaken using the Q5 High-Fidelity Polymerase kit (NEB) and thermocycling conditions of 98°C, 3 min, 35 cycles of 95°C, 30s, 55°C, 30s, 72°C, 75s. Products were visualized in ethidium-stained agarose gels following electrophoresis and recovered as above. This step was followed by an additional PCR to add homology arms (HA) to flank the reporter transgene using Gibson assembly (123). Primers were designed with HAs of 20 to 26 bp at both the promoter and the terminator sequences surrounding the EGFP coding sequence in the pJC53.2 backbone. Amplification of the promoter sequence was carried out using primers for Gibson assembly (Supplementary Table S4). A two-step PCR protocol was employed, in which the template DNA was first denatured at 98°C for 5 min, followed by 10 cycles of 95°C for 30s, 54°C for 30s, 72°C for 75s, before 25 cycles followed with 62°C for 30s, and 5 min at 72°C. The pJC53.2_SmUbi-EGFP-SmUbi construct was assembled using the NEBuilder Hifi DNA Assembly Cloning Kit (NEB). Competent *Escherichia coli* DH5α (NEB) were transformed with the ligation products, plated on LB agar supplemented with kanamycin and ampicillin (Sigma Aldrich, Darmstadt, Germany) for 18 h at 37°C, and plasmids with correct insert confirmed by Sanger sequencing (Microsynth SeqLab, Göttingen, Germany).

### Particle bombardment

Particle bombardment (PB) was carried out using the stationary PDS 1000/He system (Bio-Rad, Hercules, CA) (58, 124) to assess the activity of the SmUbi-EGFP-SmUbi transgene construct (Figure 1D). Plasmid DNA was isolated by the PureLink Expi Endotoxin-Free Maxi Plasmid Purification Kit (Thermo Fisher Scientific, Waltham, MA), and precipitated onto gold particles (0.6 µm) using CaCl_2_, spermidine, and ethanol (124); 5 µg DNA/600 ng gold. Most of the culture medium was removed, and male worms were positioned centrally in a petri dish; 10 male schistosomes were included in each biological replicate (n=5). The ballistic parameters were helium gas at 1,550 psi, 3 cm target distance (platform stage 1), 15 in Hg (381 mm Hg) atmosphere, and 23°C. After PB, the worms were immersed in culture medium to recover, and maintained for 48 h before downstream analysis. For microscopic analyses, schistosome nuclei of the worms were counterstained with Hoechst 33342 (Invitrogen, Waltham, MA) for 120 min before fluorescence microscopy (IX71 Inverted Fluorescence Microscope Pred IX73, Olympus) and confocal laser scanning microscopy (CLSM; TCS SP5 vis confocal laser scanning microscope, Leica Microsystems GmbH, Wetzlar, Germany). Captured images were analyzed with the cellSens software (Olympus) and LAS X software (Leica), respectively. Worms exhibiting EGFP fluorescence were washed in 1× PBS, transferred to 50 μl DNA/RNA protection buffer (Monarch Total RNA Miniprep Kit, NEB), and stored in liquid N_2_. Total RNA was isolated from worms, and its concentration and integrity assessed on electropherograms (RNA 6000 Nano Chips, Bioanalyzer 2100 System, Agilent Technologies, Santa Clara, CA). Reverse transcription of 100 ng total RNA from each sample to cDNA was accomplished with the QuantiTect Reverse Transcription Kit (Qiagen) at 42°C, 30 min, and the reaction terminated at 95°C, 3 min. PCRs using these cDNAs were performed in 25 µl reaction volumes with 200 μM dNTPs, 0.5 μM each primer, 1 µl cDNA, and FIREPol® DNA Polymerase (Solis BioDyne, Tartu, Estonia). The EGFP transcript of 717 nt was amplified (Figure 1E) using the primers EGFP_fw and EGFP_rev (Supplementary Table S4), and thermocycling at 98°C, 30s, 35 cycles of 98°C for 30s, 55°C for 30s, and 72°C for 60s. Ten ng pJC53.2_ SmUbi-EGFP-SmUbi plasmid-DNA served as a positive PCR control.

### Ribonucleoprotein complexes

To prepare CRISPR/Cas-mediated editing of the AT-rich GSH1 (66% AT), target protospacer adjacent motifs (PAM) and adjacent sequences for both Cas9 and Cas12a were predicted using CHOPCHOP (65, 125, 126), applying the default function against the *S. mansoni* genome. To compare the CRISPR/Cas efficiency between these two programmable endonucleases in the absence of cleavage site-specific bias, we identified guide RNA (gRNA) sites for Cas9 and Cas12a predicted to exhibit similar CRISPR efficiency (∼50%), absence of off-targets, and notably, programmed cleavages at near identical target in GSH1. Both Cas9 and Cas12a CRISPR/Cas target sites without any predicted-off-targets on the *S. mansoni* genome were used (Figure 2A). We purchased the customer-defined single CRISPR RNA (crRNA) of Cas9; sgRNA with both crRNA and tracrRNA and crRNA of Cas12a modified chemically to protect against degradation by cellular RNases (Alt-R^TM^ system, IDT, Coralville, IA). The customer-defined crRNA sequences for Cas9 and Cas12a were 5’-GAGAUCAAUUAUCUGACAAU-3’ and 5’-GAGAUCAAUUAUCUGACAAUGGAA-3’, respectively (Supplemental Table S4) (127, 128).

The Cas9-sgRNA and Cas12a-crRNA were labeled with CX-rhodamine (Label IT® Nucleic Acid Labeling Reagents Rhodamine Kit, Mirus Bio, Madison, WI) (75, 121), after which RNPs of Cas12a were assembled by mixing of 1:1 (w/w, gRNA:enzyme) Alt-R AsCas12a V3 with purified, labeled and purified crRNA; and Alt-R™ SpCas9 Nuclease V3 with labeled, purified sgRNA (121). Ten µg Cas9 and 10 µg sgRNA were mixed by pipetting in a final volume of 100 µl Opti-MEM (Gibco, Thermo Fisher) and incubated 15 min, at 23°C in the dark. Before transfection by RNPs, ∼10,000 eggs were exposed to 0.25% trypsin in EDTA at 37°C for 5 min, aiming to remove the protein lining the pores of the egg (92, 129), washed twice in 1x PBS, resuspended in 100 µl Opti-MEM in an electroporation cuvette (4 mm) (Bio-Rad, Hercules, CA) and 100 µl labeled RNPs were added. These eggs were subjected square-wave electroporation (one pulse of 125 volts, 20 ms duration (Gene Pulser Xcell Electroporation System, Bio-Rad or the BTX, ECM 830 Electro Square Porator, Holliston, MA) (26, 63). Thereafter, the eggs were maintained in complete medium as above before counterstaining for 30 min in Hoechst 333420 (H3570, Invitrogen, Waltham, MA). RNP entry into the LE were monitored under using fluorescence microscopy. CX rhodamine-specific fluorescence signals were localized by CLSM at 576 nm excitation (TCS SP5 vis confocal laser scanning microscope, Leica) (121), and images processed using LAS X software (Leica). Control groups included eggs not subjected to electroporation, eggs that were electroporated in Opti-MEM without CRISPR materials (mock treatment), and eggs transfected with an irrelevant (scramble) guide RNA/RNP complex (63).

### Donor repair template

The donor DNA template consisting of 3,351 bp of Sm*ubi* promoter (2,056 bp), EGFP (717 bp) and Sm*ubi* terminator (578 bp), and flanked by HAs of 50 bp each, were prepared by PCR using 5’ C6-PEG10-modified primers with 50 bp GSH1 (Supplementary Table S4). Yu et al. proved that DNA donor containing microhomology arms of 50 bp with 5’ modifications including an amine group, a C6 linker (AmC6) and polyethylene glycol (PEG10) treatment improve the efficient gene knock-in of a EGFP reporter gene using the CRISPR/Cas9 system (60). Here, we prepared a DNA donor template by PCR using primers specific to Sm*ubi* promoter and terminator including 50 nt of HA (Supplementary Table S4), which 5’ends were PEG10-AmC6-modified (60, 61). Briefly, 10 µM of 5’amine C6 primers (IDT, Coralville, IA) were incubated with 1 mM Bis-PEG10-NHS ester (BroadPharm, San Diego, CA) in borate buffer (Thermo Fisher) at 23°C for 18 h. Primers were desalted and dissolved in Tris-buffer using Bio-Spin 30 column (Bio-Rad) for use in PCR. The PEG10-5’AmC6-modified primers were used for the amplification of the donor template using GoTaq^®^ Green Master Mix DNA polymerase (Promega). The SmUbi-EGFP-SmUbi construct with the 600 bp HA sequence (21) targeting GSH1 served as template DNA, with thermocycling of 98°C, 5 min, 6 cycles of 95°C, 20 s, 56°C 30 s, and 72°C 3:35 min, which was followed by 25 cycles of 95°C, 20s, 68°C 3:35 min, and 72°C 3:35 min. The presence of an amplicon of 3,451 bp in length, the expected size, was confirmed following agarose gel electrophoresis, and isolated as described above.

### Programmed editing at GSH1

Genomic DNAs were recovered from LE as described (21, 26). In brief, the eggs were homogenized in DNAzol® (Molecular Research Center, Inc., Cincinnati, OH) using a motorized pestle (Kimble Chase, Thermo Fisher or PowerMasher II Nippi, Funakoshi, Tokyo, Japan), and concentration and purity assessed using a spectrophotometer. The PCR fragment flanking the expected CRISPR/Cas cut site on target GSH1 locus (∼808 bp) was amplified by GSH1 specific primers (Supplementary Table S4) with thermocycling of 98°C for 3 min, 35 cycles of 95°C for 20s, 60°C for 30s and 72°C for 60s. Amplicons were sized in gels (as above) and bands of interest isolated using the NucleoSpin Gel and PCR Clean-up kit (Takara Bio Inc., San Jose, CA). Sanger Direct Sequencing was performed at Azenta Life Sciences (South Plainfield, NJ) or Microsynth Seqlab (Göttingen, Germany). The CRISPR efficiency catalyzed by the Cas9 and Cas12a nucleases was estimated by Tracking of InDels by Decomposition (TIDE) (62) comparing with the control groups (mock and/or wild type).

To reveal the positions and frequencies of insertion-deletions (InDel) around the target GSH1 site, a fragment-harboring target GSH1 CRISPR/Cas cleavage site was amplified by specific primers containing partial Illumina adaptor sequences (Supplementary Table S4). Generated amplicons were suitable for the AmEZ service for next generation sequencing (NGS) (Azenta Life Sciences). PCR was carried out by following thermal cycling program: 98°C 3 min, 10 cycles of 95°C 20s, 55°C 30s, 72°C for 35s, and 25 cycles of 95°C 20s, 62°C 30s, and 72°C 35s. The profile of InDels at and proximal to the programmed cleavage site in GSH1 was revealed using >100,000 NGS paired reads using the CRISPREssoV2, RGEN Tool, and Cas-Analyzer (64, 66). The CRISPResso2 analysis was evaluated using NGS reads as paired end reads, using the optimal parameters: minimum homology for alignment on amplicon, 60%, base editing target base C and result base T, quantification window size to 5. The quantification window was set to –3, and the plot window size to 30. The TruSeq2PE adapter-trimming setting was used. With CRISPR RGEN, both sense and antisense reads of the sequence reads were used. In addition, the RGEN Tool was used to investigate the nuclease detection mode was set to single nuclease. All data were tested against the wild type reference, the comparison range was set to 70, the wild type marker was set to 5, and minimum frequency was set to 1.

### Transgene integration

After electroporation, LE of experimental or control groups were harvested at 5– and 10-days post transfection. Genomic DNA (gDNA) of each sample was analyzed by PCR for 5’– and 3’ transgene integration into the GSH1. To check the 5’terminus, we used a primer pair spanning 799 nt of the upstream region of GSH1 and part of the ubiquitin promoter (Figures 3B, 3C; Supplementary Table S4). For the 3’terminus, primers spanning 1,290 nt of the targeted mutation, specifically a chimeric primer spanning the EGFP coding sequence, and the terminator paired with a primer for the GSH1were used (Supplementary Table S4). For KI investigation, we designed specific primers amplifying the GSH1 located in a significant distance away from HAs of the donor DNA template combined. These primers were each combined with primers specific for the donor DNA to investigate the success of transgene integration (Figure. 3B, additional information also found in Supplementary Table S4). Both PCRs were performed by using the GoTaq^®^ Master Mix DNA polymerase (Promega, Madison, WI), 10 ng of gDNA and in a 10 µl reaction volume, as follows. PCR conditions were 98°C 5 min, 38 cycles of 95°C 30 s, 60°C 30 s, and 72°C, 80 s. Ten biological replicates were examined. The internal control for the PCR was performed by amplifying a 1,466 bp fragment using the GSH1 specific primers (Figure 3B, see also Supplementary Table S4).

The PCR conditions did not allow amplification of the 4,732 bp fragment representing the complete reporter gene integrated into GSH1. Quantitative PCR (qPCR) was performed by analyzing 10 ng gDNA (in triplicate) in 10 µL total volume of SYBR green PCR master mix (Bio-Rad, Hercules, CA) using the Bio-Rad real-time system using 400 nM of EGFP-specific primers (Supplementary Table S4). The pJC53.2_SmUbi-EGFP-SmUbi plasmid served as template for qPCR to generate an external and logarithmic standard curve based on the EGFP copy numbers, which ranged from 2.7 to 1.17×10^7^ copies. Copy number of integrated EGFP sequences were estimated using qPCR values obtained from 100 ng gDNA from eggs and interpolation from the standard curve (26, 130).

### Transgene integration analysis

EGFP expression was assessed using CLSM (Zeiss Examiner.Z1 AX10; Camera, Carl Zeiss AxioCam MRc) with an inverse objective at 20× magnification (Zeiss W Plan-Apochromat 20x/1.0 DIC (UV) VIS-IR 421402-9800), an approach that enabled imaging of the eggs in culture (20% FBS in DMEM) at 509 nm excitation (21). Images were analyzed with the ZEN 2011 (Zeiss) at days 5 and 10 following transfection using ∼3,000 LE per replicate.

Subsequently, either egg hatching was induced, or eggs were lysed for recovery of DNA and RNA (above). Transcripts of the transgene were amplified by RT-PCR. At day 10 following transfection, 3,000 LEs of each biological replicate were transferred to 50 µl RNAzol RT (Molecular Research Center, Inc., Cincinnati, OH), homogenized using a motorized pestle, and RNA isolated and quantified. Synthesis of cDNA from 30 µg total RNA per sample was carried out using the Maxima First Strand cDNA Synthesis Kit with DNase I (Thermo Fisher) and employed as template to confirm the presence of transgene transcripts using EGFP-specific primers. The SmGAPDH gene served as the reference control (Figure 3F, Supplementary Table S4) and 10 ng donor dsDNA as the positive control. PCR reaction volumes of 25 µl, with 3 µl cDNA, 1 µM each primer and GoTaq polymerase reagent (Promega, Madison, WI, USA) were subjected to thermal cycling, 98°C 3 min, 40 cycles of 95°C 30 s, 55°C 30 s, 72°C 60 s, and products assessed as above.

## DATA AVAILABILITY

Nucleotide sequence reads are available at the NIH Sequence Read Archive, BioProject PRJNA950942, accessions SRR24034375 to 24035409. All other data are provided in the text and the supplementary information.

## AUTHOR CONTRIBUTIONS

Max F. Moescheid: Methodology, Formal Analysis, Validation, Writing-original draft, Writing-review & editing. Prapakorn Wisitpongpun: Methodology, Formal Analysis. Victoria H. Mann: Methodology. Thomas Quack: Methodology. Christoph Grunau: GSH1 identification, Writing-review & editing. Christoph G. Grevelding: Conceptualization, Supervision, Writing-review & editing. Wannaporn Ittiprasert: Conceptualization, Supervision, Formal Analysis, Validation, Writing-original draft, Writing-review & editing. Paul J. Brindley: Conceptualization, Supervision, Writing-review & editing.

## ACKNOWLEDGEMENTS

1. *S. mansoni*-infected mice were provided by the Schistosomiasis Resource Center of Biomedical Research Institute, Rockville, MD through NIH-NIAID contract HHSN272201700014I for distribution through BEI Resources. The schistosome lifecycle at the Institute of Parasitology in Giessen was maintained by Christina Scheld and Georgette Stovall (Institute of Parasitology, Biomedical Research Center Seltersberg (BFS), Justus Liebig University Giessen). We thank Oliver Puckelwaldt (Institute of Parasitology, Justus Liebig University Giessen, Germany) for bioinformatical support analyzing the schistosomal Kozak-sequences. We thank Dr. James J. Collins for kindly supplying the EGFP reporter gene fragment. The scholarship of Max M0escheid was covered by in part by the Wellcome Trust FUGI grant 107475/Z/15/Z, the International Giessen Graduate Centre for the Life Sciences (GGL) of the Justus Liebig University Giessen and the German Society of Parasitology. For open access, the author applied a CC by public copyright license to any author-accepted manuscript version arising from this submission. This study also is set within the framework of the Laboratoire d’Excellence (LabEx) TULIP (ANR-10-LABX-41) with the support of LabEx CeMEB, an ANR Investments d’avenir program (ANR-10-LABX-04-01) and the Environmental Epigenomics Core Service at IHPE (Interactions Hôtes Pathogènes Environnement). Award number RR025565 (PI, Anastas Popratiloff) from the NIH Office of Research Infrastructure’s S10 program enabled the purchase of the Zeiss 710 confocal microscope.

## SUPPLEMENTARY INFORMATION

**Supplementary Figure S1.**
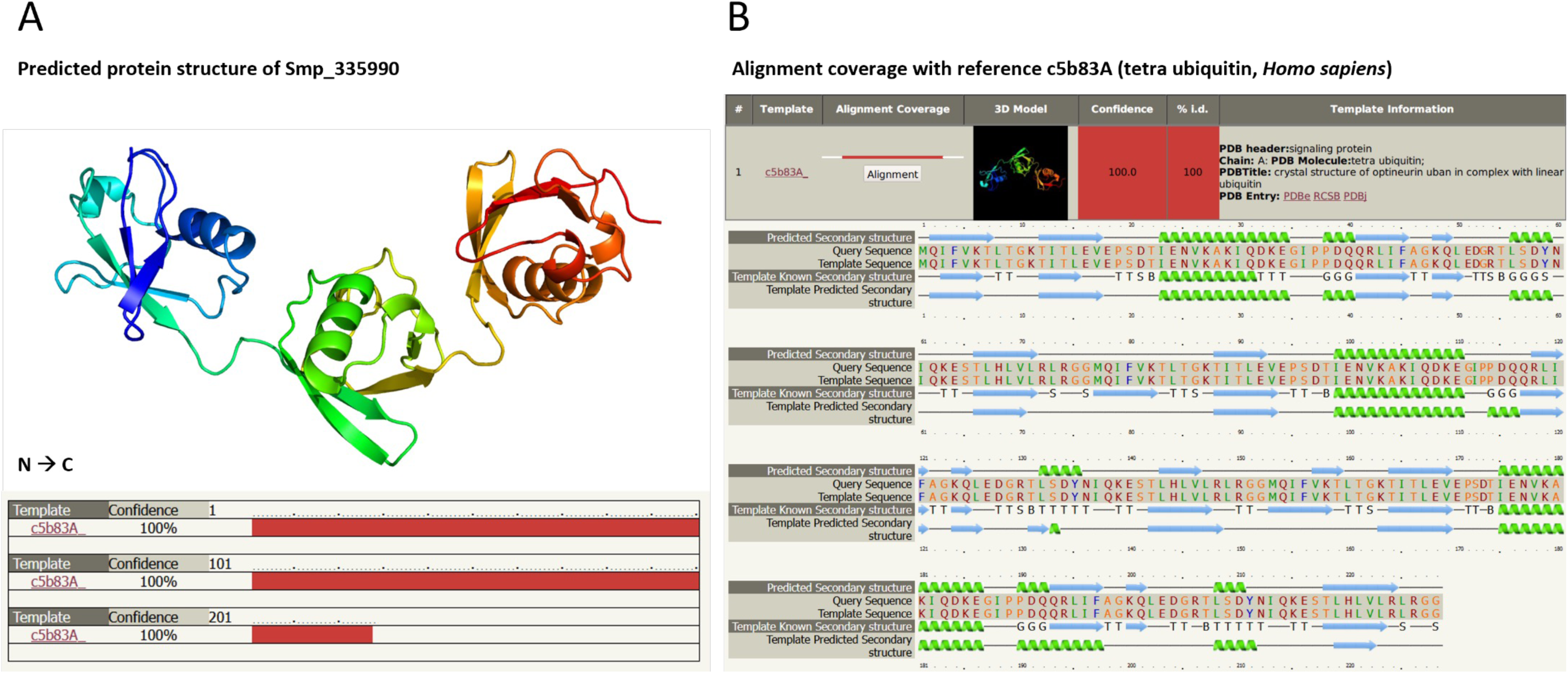
Protein structure modeling and sequence analysis confirmed Smp_335990 (SmUbi) as an orthologue of human ubiquitin. **(A)** The protein structure of SmUbi was calculated by Phyre2 (https://doi.org/10.1016/j.retram.2022.103333) based on previously known structures of orthologous ubiquitin proteins (39, 40). **(B)** The amino acid sequence of SmUbi was compared to tetra-ubiquitin orthologues from model organisms, including *Homo sapiens*, using Phyre2. SmUbi is an orthologue of human ubiquitin (UBB).

**Supplementary Figure S2.**
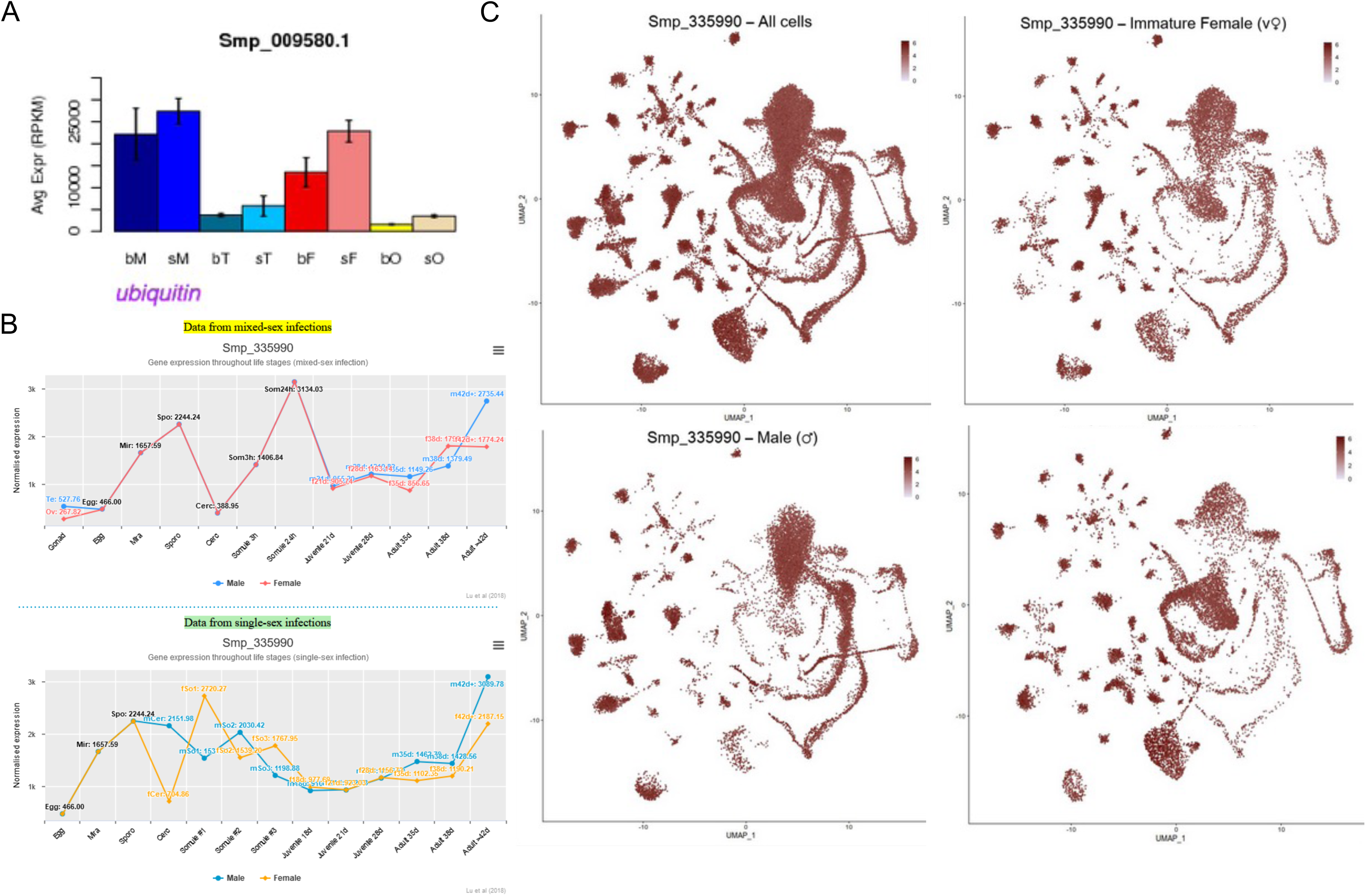
Transcriptomics data for Smp_335990 (Smubi) revealed its ubiquitous transcription patterns in both sexes and all cell types of adults and in developmental stages of *S. mansoni.* Compilation of expression data for Smp_335990 (Smubi) based on RNAseq data sets that comprised **(Panel (A))** transcript levels in adult schistosomes and their gonads (6). **(B)** The transcript levels across all life stages of (paired (top) and unpaired (bottom) *S. mansoni*, and the cell type-independent transcript profile of Smp_335990 as disclosed in the single-cell atlas of *S. mansoni* (8). Abbreviations: bM, bisex male (pairing experienced male); sM, single-sex males (pairing unexperienced male), bT, testes of bM; sT, testes of sM; bF, bisex female (pairing experienced female); sF, single-sex (pairing unexperienced female); bO, ovary of bF; sO, ovary of sF.

**Supplementary Figure S3.**
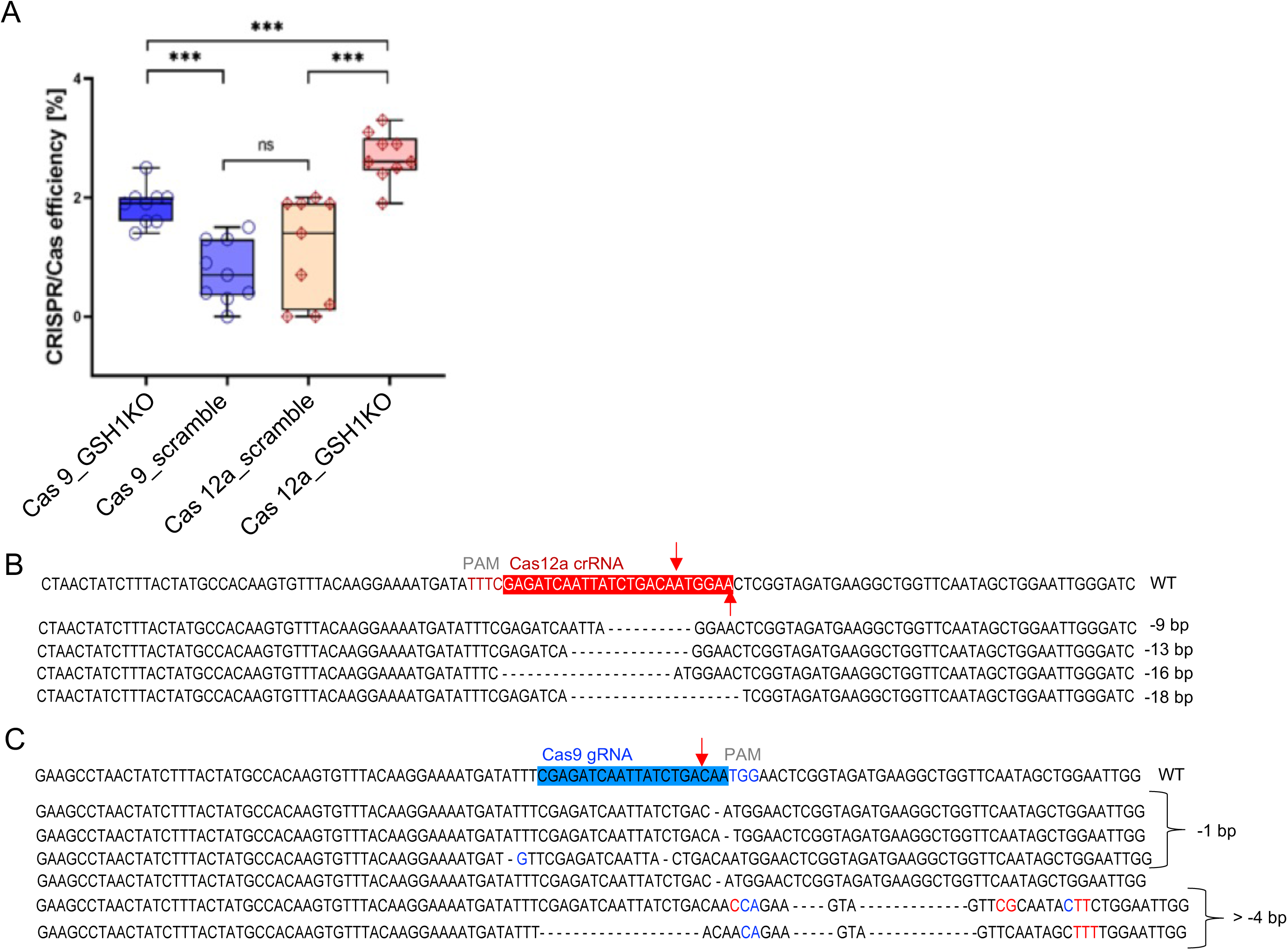
Insertions and deletions (InDels) after programmed editing of *S. mansoni* GSH1. **(A)** CRISPR/Cas efficiencies of Cas9– and Cas12a-mediated KO as determined by TIDE-analysis of adult schistosome DNA after normalization with sequencing results from the treated and control (untreated) worms. Gene editing efficiencies of the LEs, KO-Cas9 (dark blue bar), scramble-Cas9 (purple bar), scramble-Cas12a (orange bar), and KO-Cas12a (red bar) groups are presented. The programmed CRISPR mutation efficiency was significantly higher in the GSH-targeted groups than in the scramble groups (*P < 0.05, **P < 0.01, ***P < 0.001; *t* test, *n* = 8). Finally, Cas12a RNPs (red) showed significantly higher CRISPR efficiency than Cas9 RNPs. **(Panels (B) and (C))** Summary of the major InDels found at the programmed cleavage sites (red arrows) of Cas12a and Cas9, respectively, based on deep sequencing reads from three biological replicates. Abbreviations: CTRL, control; KO, knockout; ns, not significant

**Supplementary Figure S4.**
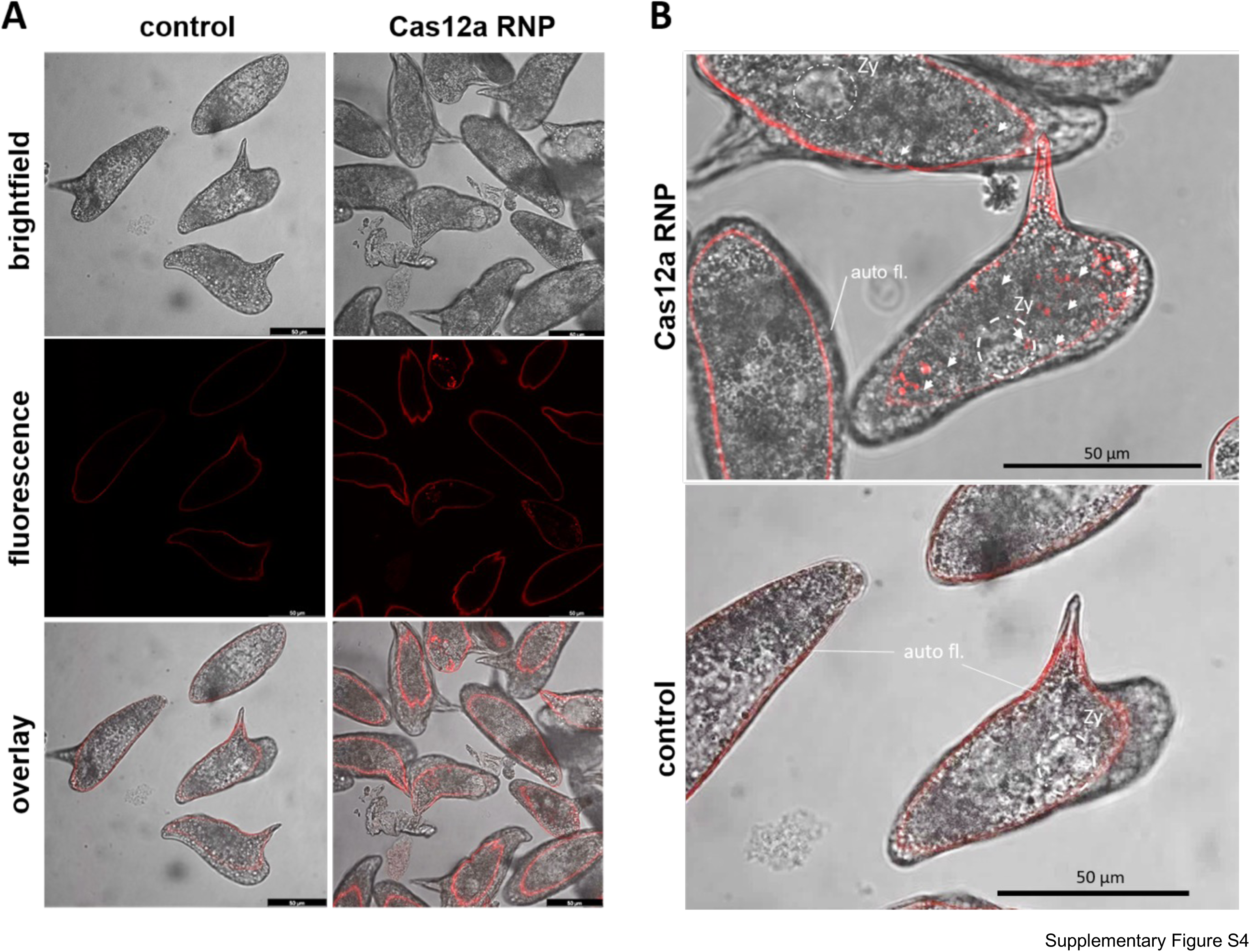
Transfection of newly laid eggs (NLE, 0 – 24 hours) with labeled CRISPR/Cas12a RNPs. **(A)** CLSM micrographs of NLE after transfection with RNPs that included rhodamine-labeled Cas12a-gRNAs. Control NLEs were subjected to electroporation in the absence of RNPs. Autofluorescence at or proximal to the eggshell was evident in the control group. By contrast, in the rhodamine labeled RNPs transfected group, red fluorescence was evident in cells of the developing larva within the egg as well as at the eggshell of many of the eggs. **(B)** Micrograph of NLEs transfected with rhodamine-labelled Cas12a-RNPs at 60 min after electroporation. Delivery of CRISPR/Cas-material (arrows) into numerous cells of the eggs was apparent. Fluorescence spectra were collected using a Leica TCS SP5 vis confocal laser scanning microscope with Leica LasX software. Scale bars: 50 µm. Abbreviations: auto fl., auto fluorescence; CLSM, confocal laser scanning microscopy; Ctr, control; Zy, Zygote.

**Supplementary Figure S5.**
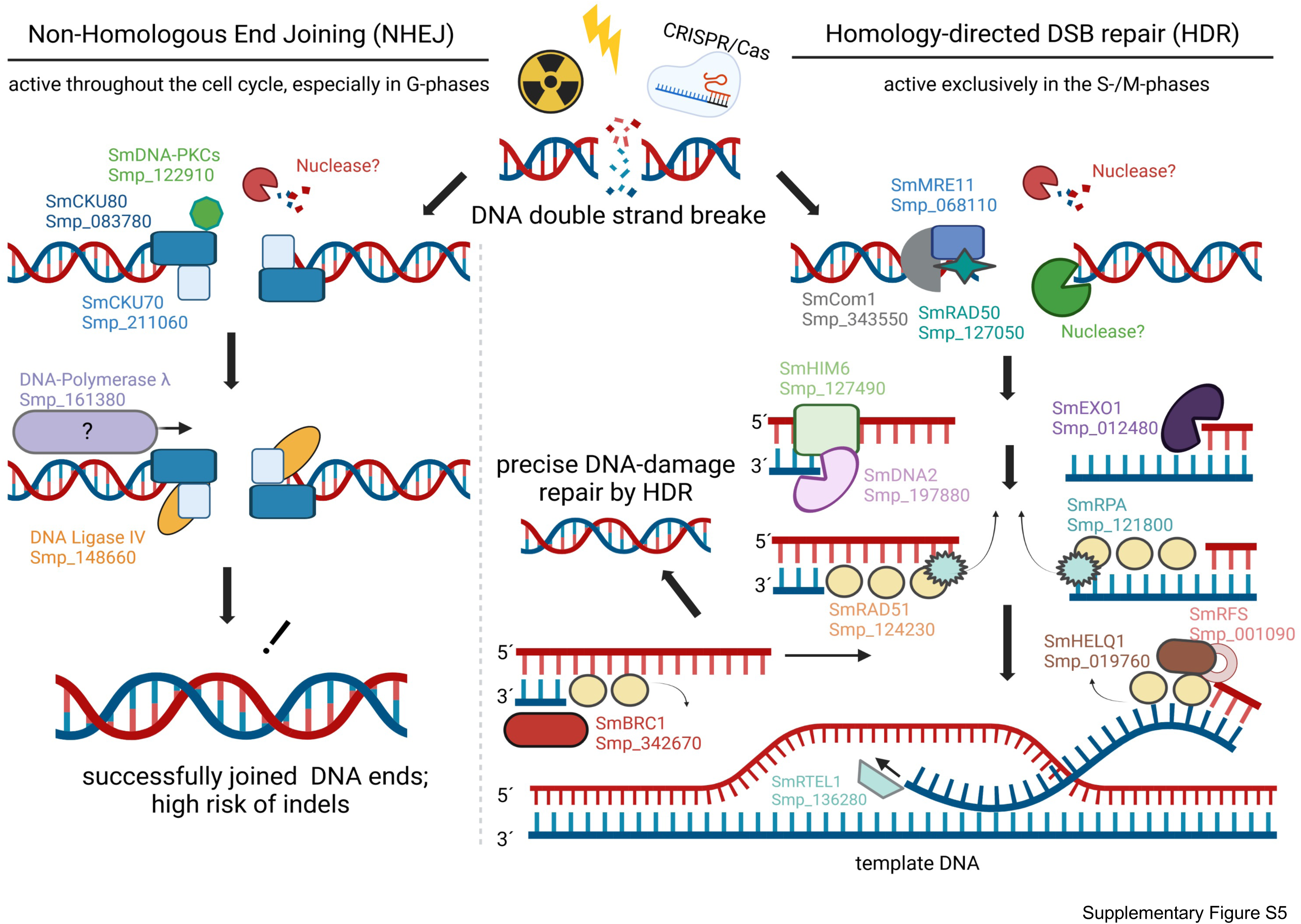
Predicted DSB-repair pathways in *S. mansoni.* Bioinformatic identification of the DNA repair machinery and cognate genes including *S. mansoni* orthologues, Smp_335990 (37, 119), as described for several model nematodes. Orthologues of the key actors of the double-stranded break (DSB)-repair pathway, as described for *Caenorhabditis elegans* (98, 99) were identified in *S. mansoni* including orthologues essential for NHEJ and for HDR. Protein sequences of DSB repair proteins from *Caenorhabditis elegans* and *Pristionchus pacificus* were analyzed using BLAST against the proteome of *S. mansoni* (parasite.wormbase.org, genome V10). Sequence coverage and domain structures were compared using the SMART Domain analyzer [smart.embl-heidelberg.de), and orthologues annotated as described (131). See also Supplementary Table S5.

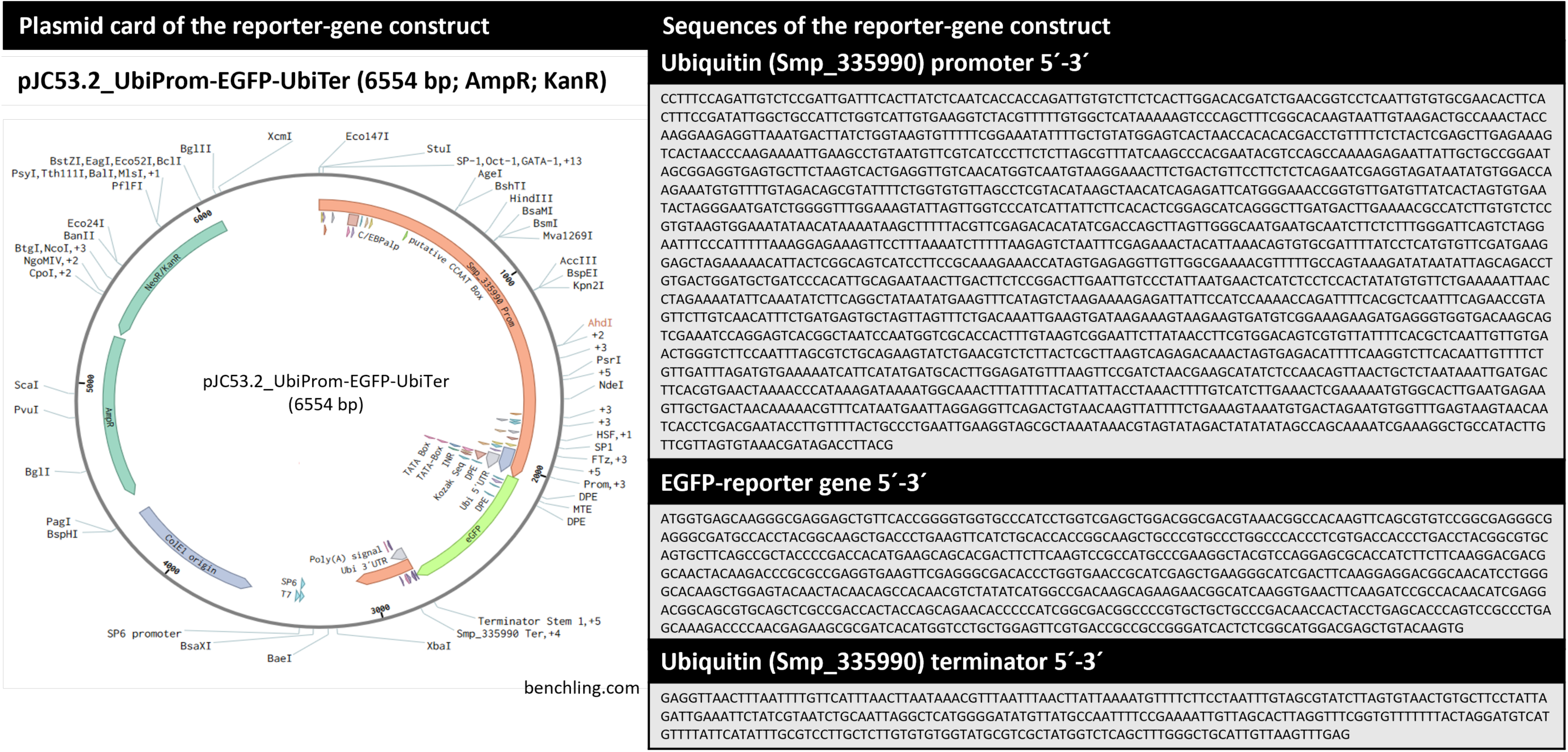
Supplementary Table 1.

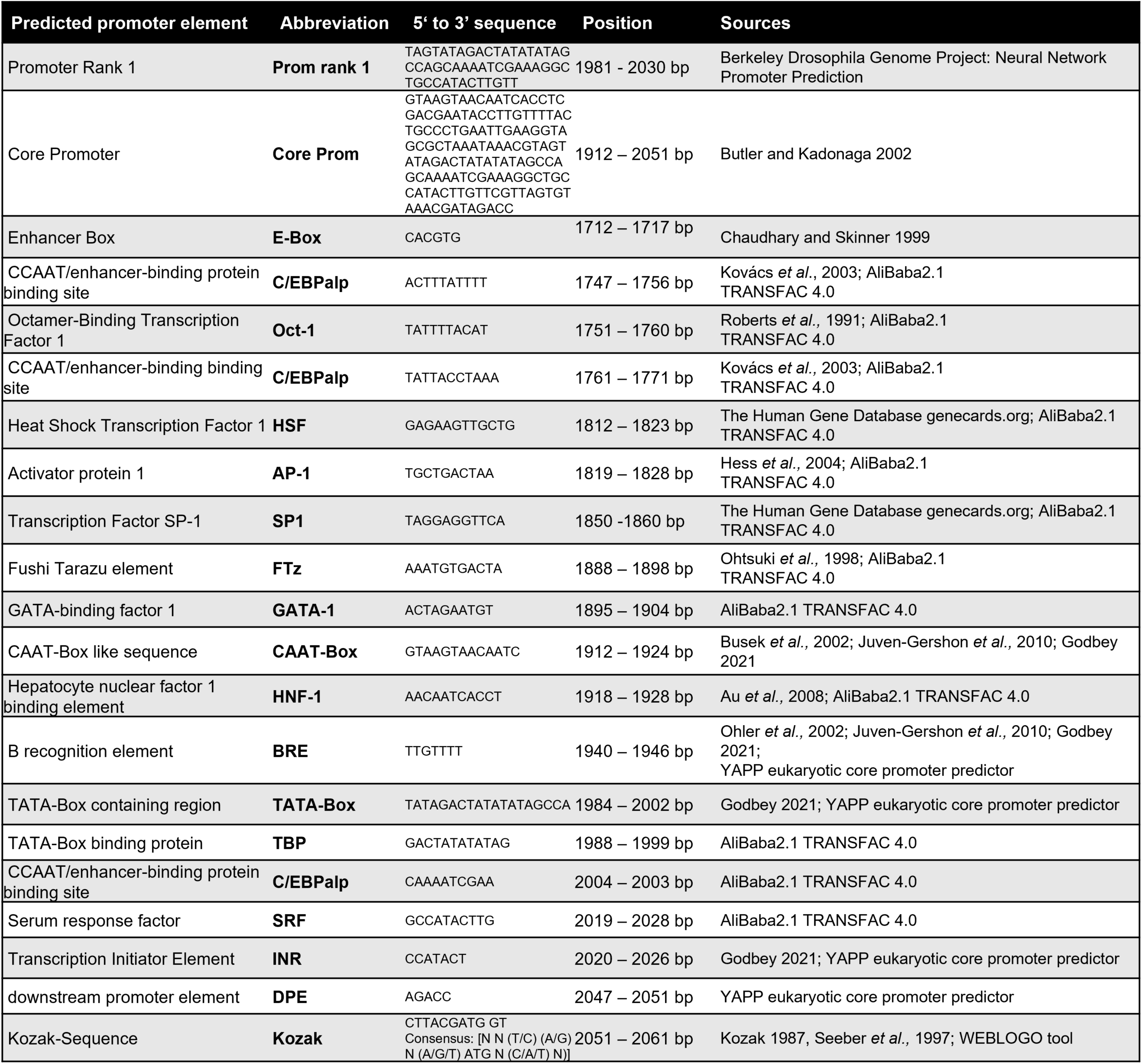
Supplementary Table 2.

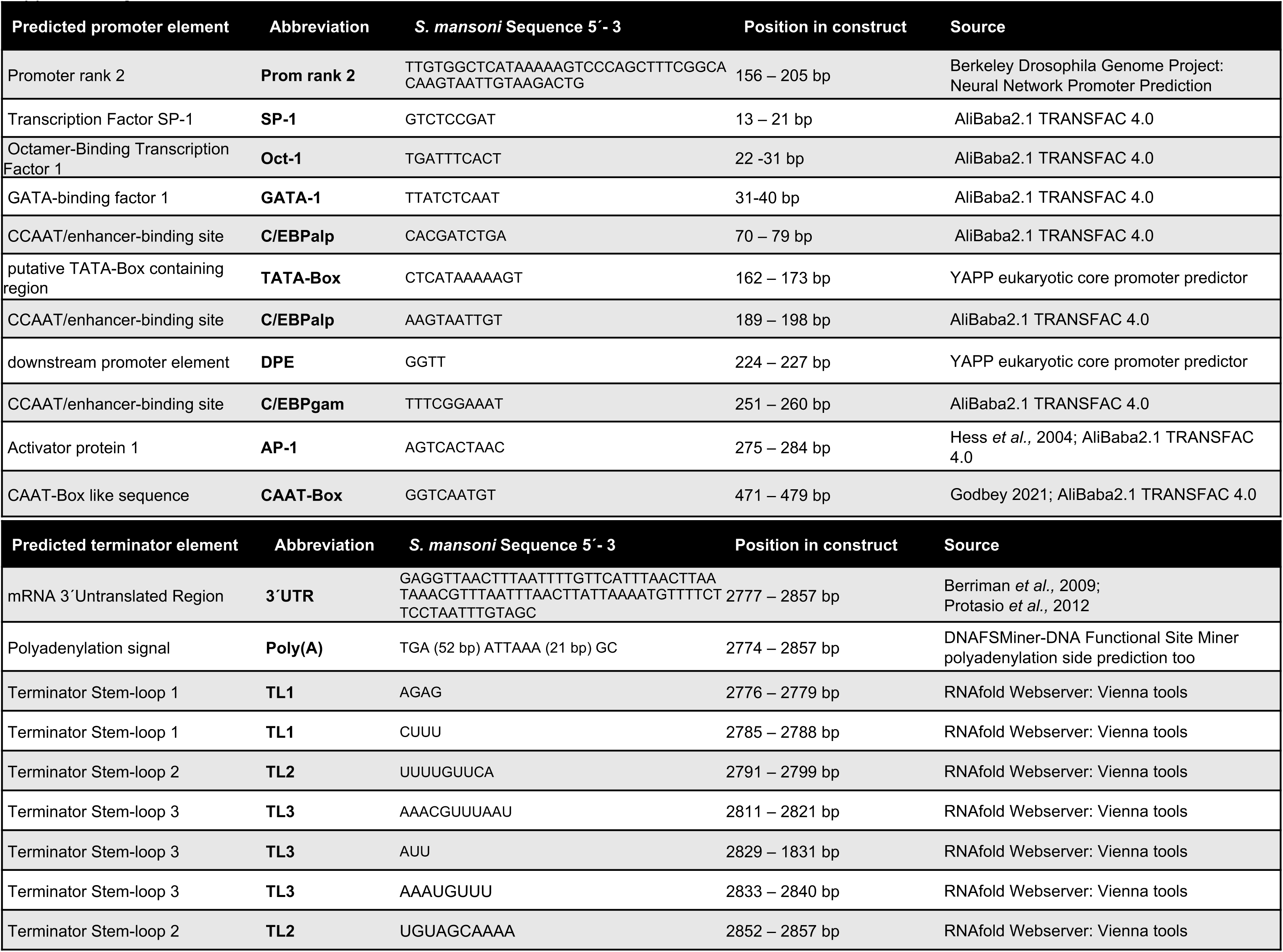
Supplementary Table 3.

**Supplemental Table S4.**
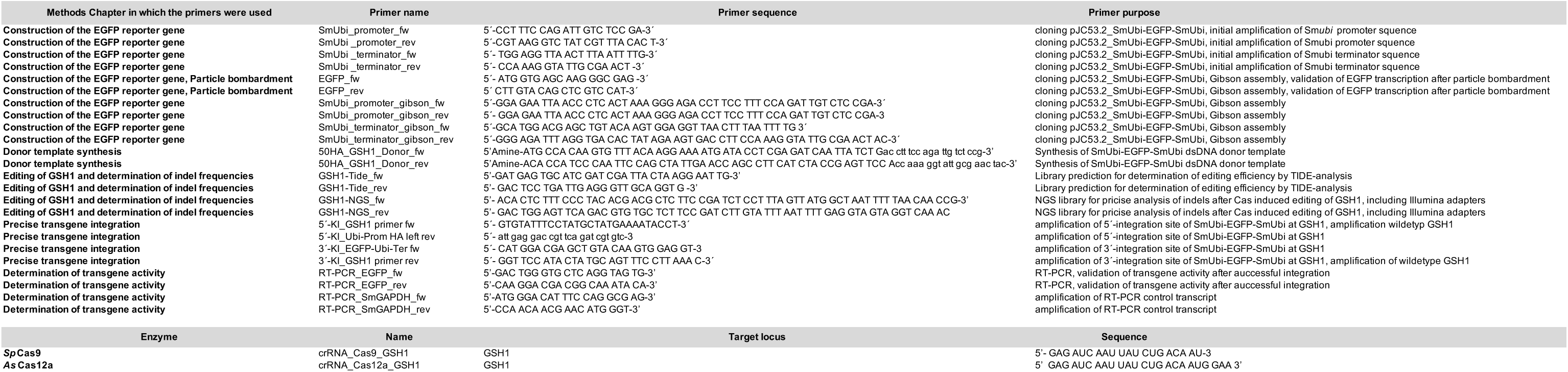
Compilation of primers used for cloning procedure, PCR, RT-PCR analysis amd crRNAs.

**Supplemental Table S5.**
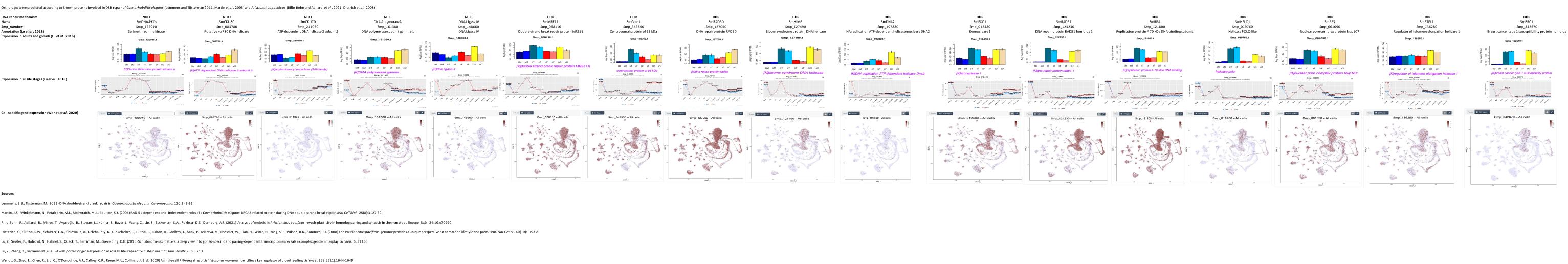
Annotation and transcriptional profiles of predicted S. mansoni double-strand break repair machinery (NHEJ and HDR).

## REFERENCES

1. Vale N, Gouveia MJ, Rinaldi G, Brindley PJ, Gartner F, Correia da Costa JM. Praziquantel for Schistosomiasis: Single-Drug Metabolism Revisited, Mode of Action, and Resistance. Antimicrob Agents Chemother. 2017;61(5).

2. Wang W, Wang L, Liang YS. Susceptibility or resistance of praziquantel in human schistosomiasis: a review. Parasitol Res. 2012;111(5):1871–7.

3. Berriman M, Haas BJ, LoVerde PT, Wilson RA, Dillon GP, Cerqueira GC, et al. The genome of the blood fluke *Schistosoma mansoni*. Nature. 2009;460(7253):352-8.

4. Crellen T, Allan F, David S, Durrant C, Huckvale T, Holroyd N, et al. Whole genome resequencing of the human parasite *Schistosoma mansoni* reveals population history and effects of selection. Sci Rep. 2016;6:20954.

5. Protasio AV, Tsai IJ, Babbage A, Nichol S, Hunt M, Aslett MA, et al. A systematically improved high quality genome and transcriptome of the human blood fluke *Schistosoma mansoni*. PLoS Negl Trop Dis. 2012;6(1):e1455.

6. Lu Z, Sessler F, Holroyd N, Hahnel S, Quack T, Berriman M, et al. Schistosome sex matters: a deep view into gonad-specific and pairing-dependent transcriptomes reveals a complex gender interplay. Sci Rep. 2016;6:31150.

7. Lu Z, Sessler F, Holroyd N, Hahnel S, Quack T, Berriman M, et al. A gene expression atlas of adult *Schistosoma mansoni* and their gonads. Sci Data. 2017;4:170118.

8. Wendt GR, Reese ML, Collins JJ, 3rd. SchistoCyte Atlas: A Single-Cell Transcriptome Resource for Adult Schistosomes. Trends Parasitol. 2021;37(7):585–7.

9. Diaz Soria CL, Lee J, Chong T, Coghlan A, Tracey A, Young MD, et al. Single-cell atlas of the first intra-mammalian developmental stage of the human parasite *Schistosoma mansoni*. Nat Commun. 2020;11(1):6411.

10. Lu Z, Sankaranarayanan G, Rawlinson KA, Offord V, Brindley PJ, Berriman M, et al. The Transcriptome of *Schistosoma mansoni* Developing Eggs Reveals Key Mediators in Pathogenesis and Life Cycle Propagation. Front Trop Dis. 2021;2:713123.

11. Wangwiwatsin A, Protasio AV, Wilson S, Owusu C, Holroyd NE, Sanders MJ, et al. Transcriptome of the parasitic flatworm *Schistosoma mansoni* during intra-mammalian development. PLoS Negl Trop Dis. 2020;14(5):e0007743.

12. Rawlinson KA, Reid AJ, Lu Z, Driguez P, Wawer A, Coghlan A, et al. Daily rhythms in gene expression of the human parasite *Schistosoma mansoni*. BMC Biol. 2021;19(1):255.

13. Vasconcelos EJR, da Silva LF, Pires DS, Lavezzo GM, Pereira ASA, Amaral MS, et al. The *Schistosoma mansoni* genome encodes thousands of long non-coding RNAs predicted to be functional at different parasite life-cycle stages. Sci Rep. 2017;7(1):10508.

14. Vasconcelos EJR, Mesel VC, da Silva LF, Pires DS, Lavezzo GM, Pereira ASA, et al. Atlas of *Schistosoma mansoni* long non-coding RNAs and their expression correlation to protein-coding genes. Database (Oxford). 2018;2018.

15. Anderson L, Amaral MS, Beckedorff F, Silva LF, Dazzani B, Oliveira KC, et al. Schistosoma mansoni Egg, Adult Male and Female Comparative Gene Expression Analysis and Identification of Novel Genes by RNA-Seq. PLoS Negl Trop Dis. 2015;9(12):e0004334.

16. Berger DJ, Crellen T, Lamberton PHL, Allan F, Tracey A, Noonan JD, et al. Whole-genome sequencing of *Schistosoma mansoni* reveals extensive diversity with limited selection despite mass drug administration. Nat Commun. 2021;12(1):4776.

17. Da’dara AA, Skelly PJ. Gene suppression in schistosomes using RNAi. Methods Mol Biol. 2015;1201:143–64.

18. Gava SG, Tavares NC, Salim ACM, Araujo FMG, Oliveira G, Mourao MM. *Schistosoma mansoni*: Off-target analyses using nonspecific double-stranded RNAs as control for RNAi experiments in schistosomula. Exp Parasitol. 2017;177:98–103.

19. Stefanic S, Dvorak J, Horn M, Braschi S, Sojka D, Ruelas DS, et al. RNA interference in *Schistosoma mansoni* schistosomula: selectivity, sensitivity and operation for larger-scale screening. PLoS Negl Trop Dis. 2010;4(10):e850.

20. Moescheid MF, Puckelwaldt O, Beutler M, Haeberlein S, Grevelding CG. Defining an optimal control for RNAi experiments with adult *Schistosoma mansoni*. Sci Rep. 2023;13(1):9766.

21. Ittiprasert W, Moescheid MF, Chaparro C, Mann VH, Quack T, Rodpai R, et al. Targeted insertion and reporter transgene activity at a gene safe harbor of the human blood fluke, Schistosoma mansoni. Cell Rep Methods. 2023;3(7):100535.

22. Zhang B. CRISPR/Cas gene therapy. J Cell Physiol. 2021;236(4):2459–81.

23. Zhang D, Zhang Z, Unver T, Zhang B. CRISPR/Cas: A powerful tool for gene function study and crop improvement. J Adv Res. 2021;29:207–21.

24. Makarova KS, Wolf YI, Iranzo J, Shmakov SA, Alkhnbashi OS, Brouns SJJ, et al. Evolutionary classification of CRISPR-Cas systems: a burst of class 2 and derived variants. Nat Rev Microbiol. 2020;18(2):67–83.

25. Pickar-Oliver A, Gersbach CA. The next generation of CRISPR-Cas technologies and applications. Nat Rev Mol Cell Biol. 2019;20(8):490–507.

26. Ittiprasert W, Chatupheeraphat C, Mann VH, Li W, Miller A, Ogunbayo T, et al. RNA-Guided AsCas12a– and SpCas9-Catalyzed Knockout and Homology Directed Repair of the Omega-1 Locus of the Human Blood Fluke, *Schistosoma mansoni*. Int J Mol Sci. 2022;23(2).

27. Cong L, Ran FA, Cox D, Lin S, Barretto R, Habib N, et al. Multiplex genome engineering using CRISPR/Cas systems. Science. 2013;339(6121):819-23.

28. Jinek M, Chylinski K, Fonfara I, Hauer M, Doudna JA, Charpentier E. A programmable dual-RNA-guided DNA endonuclease in adaptive bacterial immunity. Science. 2012;337(6096):816–21.

29. Mali P, Yang L, Esvelt KM, Aach J, Guell M, DiCarlo JE, et al. RNA-guided human genome engineering via Cas9. Science. 2013;339(6121):823–6.

30. Paul B, Montoya G. CRISPR-Cas12a: Functional overview and applications. Biomed J. 2020;43(1):8–17.

31. Swarts DC, Jinek M. Cas9 versus Cas12a/Cpf1: Structure-function comparisons and implications for genome editing. Wiley Interdiscip Rev RNA. 2018;9(5):e1481.

32. Kim H, Kim ST, Ryu J, Kang BC, Kim JS, Kim SG. CRISPR/Cpf1-mediated DNA-free plant genome editing. Nat Commun. 2017;8:14406.

33. Nekrasov V, Staskawicz B, Weigel D, Jones JD, Kamoun S. Targeted mutagenesis in the model plant *Nicotiana benthamiana* using Cas9 RNA-guided endonuclease. Nat Biotechnol. 2013;31(8):691–3.

34. Svitashev S, Young JK, Schwartz C, Gao H, Falco SC, Cigan AM. Targeted Mutagenesis, Precise Gene Editing, and Site-Specific Gene Insertion in Maize Using Cas9 and Guide RNA. Plant Physiol. 2015;169(2):931–45.

35. Papapetrou EP, Schambach A. Gene Insertion Into Genomic Safe Harbors for Human Gene Therapy. Mol Ther. 2016;24(4):678–84.

36. Jurberg AD, Goncalves T, Costa TA, de Mattos AC, Pascarelli BM, de Manso PP, et al. The embryonic development of *Schistosoma mansoni* eggs: proposal for a new staging system. Dev Genes Evol. 2009;219(5):219–34.

37. Howe KL, Bolt BJ, Shafie M, Kersey P, Berriman M. WormBase ParaSite – a comprehensive resource for helminth genomics. Mol Biochem Parasitol. 2017;215:2–10.

38. Kelley LA, Mezulis S, Yates CM, Wass MN, Sternberg MJ. The Phyre2 web portal for protein modeling, prediction and analysis. Nat Protoc. 2015;10(6):845–58.

39. Gao S, Pan M, Zheng Y, Huang Y, Zheng Q, Sun D, et al. Monomer/Oligomer Quasi-Racemic Protein Crystallography. J Am Chem Soc. 2016;138(43):14497–502.

40. Silver CE, Cusumano RJ, Fell SC, Strauch B. Replacement of upper esophagus: results with myocutaneous flap and with gastric transposition. Laryngoscope. 1989;99(8 Pt 1):819–21.

41. Webb GC, Baker RT, Fagan K, Board PG. Localization of the human UbB polyubiquitin gene to chromosome band 17p11.1-17p12. Am J Hum Genet. 1990;46(2):308–15.

42. Reese MG. Application of a time-delay neural network to promoter annotation in the *Drosophila melanogaster* genome. Comput Chem. 2001;26(1):51–6.

43. Roberts SB, Segil N, Heintz N. Differential phosphorylation of the transcription factor Oct1 during the cell cycle. Science. 1991;253(5023):1022–6.

44. Wingender E. TRANSFAC, TRANSPATH and CYTOMER as starting points for an ontology of regulatory networks. In Silico Biol. 2004;4(1):55–61.

45. Gershenzon NI, Ioshikhes IP. Synergy of human Pol II core promoter elements revealed by statistical sequence analysis. Bioinformatics. 2005;21(8):1295–300.

46. Jin VX, Singer GA, Agosto-Perez FJ, Liyanarachchi S, Davuluri RV. Genome-wide analysis of core promoter elements from conserved human and mouse orthologous pairs. BMC Bioinformatics. 2006;7:114.

47. Chalkley GE, Verrijzer CP. DNA binding site selection by RNA polymerase II TAFs: a TAF(II)250-TAF(II)150 complex recognizes the initiator. EMBO J. 1999;18(17):4835–45.

48. Cartharius K, Frech K, Grote K, Klocke B, Haltmeier M, Klingenhoff A, et al. MatInspector and beyond: promoter analysis based on transcription factor binding sites. Bioinformatics. 2005;21(13):2933–42.

49. Butler JE, Kadonaga JT. The RNA polymerase II core promoter: a key component in the regulation of gene expression. Genes Dev. 2002;16(20):2583–92.

50. Kovacs KA, Steinmann M, Magistretti PJ, Halfon O, Cardinaux JR. CCAAT/enhancer-binding protein family members recruit the coactivator CREB-binding protein and trigger its phosphorylation. J Biol Chem. 2003;278(38):36959–65.

51. Godbey WT. Biotechnology and its Applications. 2nd ed: Elsevier; 2021.

52. Kozak M. At least six nucleotides preceding the AUG initiator codon enhance translation in mammalian cells. J Mol Biol. 1987;196(4):947–50.

53. Lu Z, Zhang Y, Berriman M. A web portal for gene expression across all life stages of Schistosoma mansoni. 2018.

54. Wendt G, Zhao L, Chen R, Liu C, O’Donoghue AJ, Caffrey CR, et al. A single-cell RNA-seq atlas of *Schistosoma mansoni* identifies a key regulator of blood feeding. Science. 2020;369(6511):1644–9.

55. Liu H, Han H, Li J, Wong L. DNAFSMiner: a web-based software toolbox to recognize two types of functional sites in DNA sequences. Bioinformatics. 2005;21(5):671–3.

56. Gruber AR, Lorenz R, Bernhart SH, Neubock R, Hofacker IL. The Vienna RNA websuite. Nucleic Acids Res. 2008;36(Web Server issue):W70–4.

57. Cheng SW, Lynch EC, Leason KR, Court DL, Shapiro BA, Friedman DI. Functional importance of sequence in the stem-loop of a transcription terminator. Science. 1991;254(5035):1205–7.

58. Wippersteg V, Kapp K, Kunz W, Jackstadt WP, Zahner H, Grevelding CG. HSP70-controlled GFP expression in transiently transformed schistosomes. Mol Biochem Parasitol. 2002;120(1):141–50.

59. Wippersteg V, Ribeiro F, Liedtke S, Kusel JR, Grevelding CG. The uptake of Texas Red-BSA in the excretory system of schistosomes and its colocalisation with ER60 promoter-induced GFP in transiently transformed adult males. Int J Parasitol. 2003;33(11):1139–43.

60. Yu Y, Guo Y, Tian Q, Lan Y, Yeh H, Zhang M, et al. Publisher Correction: An efficient gene knock-in strategy using 5’-modified double-stranded DNA donors with short homology arms. Nat Chem Biol. 2020;16(4):479.

61. Yu Y, Guo Y, Tian Q, Lan Y, Yeh H, Zhang M, et al. An efficient gene knock-in strategy using 5’-modified double-stranded DNA donors with short homology arms. Nat Chem Biol. 2020;16(4):387–90.

62. Brinkman EK, van Steensel B. Rapid Quantitative Evaluation of CRISPR Genome Editing by TIDE and TIDER. Methods Mol Biol. 2019;1961:29–44.

63. Hulme BJ, Geyer KK, Forde-Thomas JE, Padalino G, Phillips DW, Ittiprasert W, et al. *Schistosoma mansoni* alpha-N-acetylgalactosaminidase (SmNAGAL) regulates coordinated parasite movement and egg production. PLoS Pathog. 2022;18(1):e1009828.

64. Clement K, Rees H, Canver MC, Gehrke JM, Farouni R, Hsu JY, et al. CRISPResso2 provides accurate and rapid genome editing sequence analysis. Nat Biotechnol. 2019;37(3):224–6.

65. Labun K, Montague TG, Gagnon JA, Thyme SB, Valen E. CHOPCHOP v2: a web tool for the next generation of CRISPR genome engineering. Nucleic Acids Res. 2016;44(W1):W272–6.

66. Park J, Lim K, Kim JS, Bae S. Cas-analyzer: an online tool for assessing genome editing results using NGS data. Bioinformatics. 2017;33(2):286–8.

67. Hotez PJ, Aksoy S, Brindley PJ, Kamhawi S. What constitutes a neglected tropical disease? PLoS Negl Trop Dis. 2020;14(1):e0008001.

68. Mitra AK, Mawson AR. Neglected Tropical Diseases: Epidemiology and Global Burden. Trop Med Infect Dis. 2017;2(3).

69. Tidman R, Kanankege KST, Bangert M, Abela-Ridder B. Global prevalence of 4 neglected foodborne trematodes targeted for control by WHO: A scoping review to highlight the gaps. PLoS Negl Trop Dis. 2023;17(3):e0011073.

70. Aya Pastrana N, Beran D, Somerville C, Heller O, Correia JC, Suggs LS. The process of building the priority of neglected tropical diseases: A global policy analysis. PLoS Negl Trop Dis. 2020;14(8):e0008498.

71. Ittiprasert W, Mann VH, Karinshak SE, Coghlan A, Rinaldi G, Sankaranarayanan G, et al. Programmed genome editing of the omega-1 ribonuclease of the blood fluke, *Schistosoma mansoni*. Elife. 2019;8:e41337.

72. Stitz M, Chaparro C, Lu Z, Olzog VJ, Weinberg CE, Blom J, et al. Satellite-Like W-Elements: Repetitive, Transcribed, and Putative Mobile Genetic Factors with Potential Roles for Biology and Evolution of Schistosoma mansoni. Genome Biol Evol. 2021;13(10).

73. Roquis D, Taudt A, Geyer KK, Padalino G, Hoffmann KF, Holroyd N, et al. Histone methylation changes are required for life cycle progression in the human parasite Schistosoma mansoni. PLoS Pathog. 2018;14(5):e1007066.

74. Wang X, Tang Y, Lu J, Shao Y, Qin X, Li Y, et al. Characterization of novel cytochrome P450 2E1 knockout rat model generated by CRISPR/Cas9. Biochem Pharmacol. 2016;105:80–90.

75. Sankaranarayanan G, Coghlan A, Driguez P, Lotkowska ME, Sanders M, Holroyd N, et al. Large CRISPR-Cas-induced deletions in the oxamniquine resistance locus of the human parasite *Schistosoma mansoni*. Wellcome Open Res. 2020;5:178.

76. Chaiyadet S, Tangkawattana S, Smout MJ, Ittiprasert W, Mann VH, Deenonpoe R, et al. Knockout of liver fluke granulin, Ov-grn-1, impedes malignant transformation during chronic infection with Opisthorchis viverrini. PLoS Pathog. 2022;18(9):e1010839.

77. Khampoosa P, Jones MK, Lovas EM, Srisawangwong T, Laha T, Piratae S, et al. Light and electron microscopy observations of embryogenesis and egg development in the human liver fluke, *Opisthorchis viverrini* (Platyhelminthes, Digenea). Parasitol Res. 2012;110(2):799–808.

78. Born-Torrijos A, Holzer AS, Raga JA, van Beest GS, Yoneva A. Description of embryonic development and ultrastructure in miracidia of *Cardiocephaloides longicollis* (Digenea, Strigeidae) in relation to active host finding strategy in a marine environment. J Morphol. 2017;278(8):1137–48.

79. You H, Mayer JU, Johnston RL, Sivakumaran H, Ranasinghe S, Rivera V, et al. CRISPR/Cas9-mediated genome editing of *Schistosoma mansoni* acetylcholinesterase. FASEB J. 2021;35(1):e21205.

80. Du X, McManus DP, French JD, Collinson N, Sivakumaran H, MacGregor SR, et al. CRISPR interference for sequence-specific regulation of fibroblast growth factor receptor A in *Schistosoma mansoni*. Front Immunol. 2022;13:1105719.

81. Race GJ, Michaels RM, Martin JH, Larsh JE, Jr., Matthews JL. *Schistosoma mansoni* eggs: an electron microscopic study of shell pores and microbarbs. Proc Soc Exp Biol Med. 1969;130(3):990–2.

82. Kazemian P, Yu SY, Thomson SB, Birkenshaw A, Leavitt BR, Ross CJD. Lipid-Nanoparticle-Based Delivery of CRISPR/Cas9 Genome-Editing Components. Mol Pharm. 2022;19(6):1669–86.

83. Kuppers DA, Linton J, Ortiz Espinosa S, McKenna KM, Rongvaux A, Paddison PJ. Gene knock-outs in human CD34+ hematopoietic stem and progenitor cells and in the human immune system of mice. PLoS One. 2023;18(6):e0287052.

84. Mazurov D, Ramadan L, Kruglova N. Packaging and Uncoating of CRISPR/Cas Ribonucleoproteins for Efficient Gene Editing with Viral and Non-Viral Extracellular Nanoparticles. Viruses. 2023;15(3).

85. Bhandawat A, Sharma V, Rishi V, J KR. Biolistic Delivery of Programmable Nuclease (CRISPR/Cas9) in Bread Wheat. Methods Mol Biol. 2020;2124:309–29.

86. Heyers O, Walduck AK, Brindley PJ, Bleiss W, Lucius R, Dorbic T, et al. *Schistosoma mansoni* miracidia transformed by particle bombardment infect Biomphalaria glabrata snails and develop into transgenic sporocysts. Exp Parasitol. 2003;105(2):174–8.

87. Wippersteg V, Kapp K, Kunz W, Grevelding CG. Characterisation of the cysteine protease ER60 in transgenic *Schistosoma mansoni* larvae. Int J Parasitol. 2002;32(10):1219–24.

88. Khan S, Sallard E. Current and Prospective Applications of CRISPR-Cas12a in Pluricellular Organisms. Mol Biotechnol. 2023;65(2):196–205.

89. Moreno-Mateos MA, Fernandez JP, Rouet R, Vejnar CE, Lane MA, Mis E, et al. CRISPR-Cpf1 mediates efficient homology-directed repair and temperature-controlled genome editing. Nat Commun. 2017;8(1):2024.

90. Mao Z, Bozzella M, Seluanov A, Gorbunova V. DNA repair by nonhomologous end joining and homologous recombination during cell cycle in human cells. Cell Cycle. 2008;7(18):2902–6.

91. Lu Z, Quack T, Hahnel S, Gelmedin V, Pouokam E, Diener M, et al. Isolation, enrichment and primary characterisation of vitelline cells from *Schistosoma mansoni* obtained by the organ isolation method. Int J Parasitol. 2015;45(9-10):663–72.

92. Neill PJ, Smith JH, Doughty BL, Kemp M. The ultrastructure of the *Schistosoma mansoni* egg. Am J Trop Med Hyg. 1988;39(1):52–65.

93. Ashton PD, Harrop R, Shah B, Wilson RA. The schistosome egg: development and secretions. Parasitology. 2001;122(Pt 3):329–38.

94. Fattah F, Lee EH, Weisensel N, Wang Y, Lichter N, Hendrickson EA. Ku regulates the non-homologous end joining pathway choice of DNA double-strand break repair in human somatic cells. PLoS Genet. 2010;6(2):e1000855.

95. Sato M, Takabayashi S, Akasaka E, Nakamura S. Recent Advances and Future Perspectives of In Vivo Targeted Delivery of Genome-Editing Reagents to Germ Cells, Embryos, and Fetuses in Mice. Cells. 2020;9(4).

96. Hoch NC. Tissue Specificity of DNA Damage and Repair. Physiology (Bethesda). 2023;38(5):0.

97. Iyama T, Wilson DM, 3rd. DNA repair mechanisms in dividing and non-dividing cells. DNA Repair (Amst). 2013;12(8):620–36.

98. Bae W, Hong S, Park MS, Jeong HK, Lee MH, Koo HS. Single-strand annealing mediates the conservative repair of double-strand DNA breaks in homologous recombination-defective germ cells of *Caenorhabditis elegans*. DNA Repair (Amst). 2019;75:18–28.

99. Lemmens BB, Tijsterman M. DNA double-strand break repair in *Caenorhabditis elegans*. Chromosoma. 2011;120(1):1–21.

100. Dieterich C, Clifton SW, Schuster LN, Chinwalla A, Delehaunty K, Dinkelacker I, et al. The *Pristionchus pacificus* genome provides a unique perspective on nematode lifestyle and parasitism. Nat Genet. 2008;40(10):1193–8.

101. Rillo-Bohn R, Adilardi R, Mitros T, Avsaroglu B, Stevens L, Kohler S, et al. Analysis of meiosis in *Pristionchus pacificus* reveals plasticity in homolog pairing and synapsis in the nematode lineage. Elife. 2021;10.

102. Consortium CeS. Genome sequence of the nematode *C. elegans*: a platform for investigating biology. Science. 1998;282(5396):2012–8.

103. Martin JS, Winkelmann N, Petalcorin MI, McIlwraith MJ, Boulton SJ. RAD-51-dependent and – independent roles of a *Caenorhabditis elegans* BRCA2-related protein during DNA double-strand break repair. Mol Cell Biol. 2005;25(8):3127–39.

104. Lieber MR. The mechanism of human nonhomologous DNA end joining. J Biol Chem. 2008;283(1):1–5.

105. Ozturk S, Demir N. DNA repair mechanisms in mammalian germ cells. Histol Histopathol. 2011;26(4):505–17.

106. You H, Jones MK, Whitworth DJ, McManus DP. Innovations and Advances in Schistosome Stem Cell Research. Front Immunol. 2021;12:599014.

107. Ohler U, Liao GC, Niemann H, Rubin GM. Computational analysis of core promoters in the *Drosophila* genome. Genome Biol. 2002;3(12):RESEARCH0087.

108. Juven-Gershon T, Kadonaga JT. Regulation of gene expression via the core promoter and the basal transcriptional machinery. Dev Biol. 2010;339(2):225–9.

109. Busek SU, Fantappie M, Malaquias LC, Wilson RA, Correa-Oliveira R, Oliveira GC. Cis-acting elements, CArG-, E-, CCAAT– and TATA-boxes may be involved in sexually regulated gene transcription in Schistosoma mansoni. Mem Inst Oswaldo Cruz. 2002;97 Suppl 1:85–90.

110. Byrne SM, Ortiz L, Mali P, Aach J, Church GM. Multi-kilobase homozygous targeted gene replacement in human induced pluripotent stem cells. Nucleic Acids Res. 2015;43(3):e21.

111. Zhang JP, Li XL, Li GH, Chen W, Arakaki C, Botimer GD, et al. Efficient precise knockin with a double cut HDR donor after CRISPR/Cas9-mediated double-stranded DNA cleavage. Genome Biol. 2017;18(1):35.

112. Sakuma T, Takenaga M, Kawabe Y, Nakamura T, Kamihira M, Yamamoto T. Homologous Recombination-Independent Large Gene Cassette Knock-in in CHO Cells Using TALEN and MMEJ-Directed Donor Plasmids. Int J Mol Sci. 2015;16(10):23849–66.

113. Sakuma T, Nakade S, Sakane Y, Suzuki KT, Yamamoto T. MMEJ-assisted gene knock-in using TALENs and CRISPR-Cas9 with the PITCh systems. Nat Protoc. 2016;11(1):118–33.

114. Li G, Zhang X, Wang H, Mo J, Zhong C, Shi J, et al. CRISPR/Cas9-Mediated Integration of Large Transgene into Pig CEP112 Locus. G3 (Bethesda). 2020;10(2):467–73.

115. Geldhof P, Visser A, Clark D, Saunders G, Britton C, Gilleard J, et al. RNA interference in parasitic helminths: current situation, potential pitfalls and future prospects. Parasitology. 2007;134(Pt 5):609–19.

116. Mann VH, Morales ME, Rinaldi G, Brindley PJ. Culture for genetic manipulation of developmental stages of *Schistosoma mansoni*. Parasitology. 2010;137(3):451–62.

117. Dalton JP, Clough KA, Jones MK, Brindley PJ. The cysteine proteinases of *Schistosoma mansoni* cercariae. Parasitology. 1997;114 (Pt 2):105–12.

118. Cormack BP, Valdivia RH, Falkow S. FACS-optimized mutants of the green fluorescent protein (GFP). Gene. 1996;173(1 Spec No):33–8.

119. Howe KL, Bolt BJ, Cain S, Chan J, Chen WJ, Davis P, et al. WormBase 2016: expanding to enable helminth genomic research. Nucleic Acids Res. 2016;44(D1):D774–80.

120. Crooks GE, Hon G, Chandonia JM, Brenner SE. WebLogo: a sequence logo generator. Genome Res. 2004;14(6):1188–90.

121. Nasri M, Mir P, Dannenmann B, Amend D, Skroblyn T, Xu Y, et al. Fluorescent labeling of CRISPR/Cas9 RNP for gene knockout in HSPCs and iPSCs reveals an essential role for GADD45b in stress response. Blood Adv. 2019;3(1):63–71.

122. Collins JJ, 3rd, Hou X, Romanova EV, Lambrus BG, Miller CM, Saberi A, et al. Genome-wide analyses reveal a role for peptide hormones in planarian germline development. PLoS Biol. 2010;8(10):e1000509.

123. Gibson DG, Young L, Chuang RY, Venter JC, Hutchison CA, 3rd, Smith HO. Enzymatic assembly of DNA molecules up to several hundred kilobases. Nat Methods. 2009;6(5):343–5.

124. Beckmann S, Wippersteg V, El-Bahay A, Hirzmann J, Oliveira G, Grevelding CG. *Schistosoma mansoni*: germ-line transformation approaches and actin-promoter analysis. Exp Parasitol. 2007;117(3):292–303.

125. Labun K, Montague TG, Krause M, Torres Cleuren YN, Tjeldnes H, Valen E. CHOPCHOP v3: expanding the CRISPR web toolbox beyond genome editing. Nucleic Acids Res. 2019;47(W1):W171–W4.

126. Montague TG, Cruz JM, Gagnon JA, Church GM, Valen E. CHOPCHOP: a CRISPR/Cas9 and TALEN web tool for genome editing. Nucleic Acids Res. 2014;42(Web Server issue):W401–7.

127. Jeon Y, Choi YH, Jang Y, Yu J, Goo J, Lee G, et al. Direct observation of DNA target searching and cleavage by CRISPR-Cas12a. Nat Commun. 2018;9(1):2777.

128. Zetsche B, Gootenberg JS, Abudayyeh OO, Slaymaker IM, Makarova KS, Essletzbichler P, et al. Cpf1 is a single RNA-guided endonuclease of a class 2 CRISPR-Cas system. Cell. 2015;163(3):759–71.

129. Candido RRF, Morassutti AL, Graeff-Teixeira C, St Pierre TG, Jones MK. Exploring Structural and Physical Properties of Schistosome Eggs: Potential Pathways for Novel Diagnostics? Adv Parasitol. 2018;100:209–37.

130. Joshi M, Keith Pittman H, Haisch C, Verbanac K. Real-time PCR to determine transgene copy number and to quantitate the biolocalization of adoptively transferred cells from EGFP-transgenic mice. Biotechniques. 2008;45(3):247–58.

131. Dalton JP, Day SR, Drew AC, Brindley PJ. A method for the isolation of schistosome eggs and miracidia free of contaminating host tissues. Parasitology. 1997;115 (Pt 1):29–32.

